# SLC38A2 provides proline to fulfil unique synthetic demands arising during osteoblast differentiation and bone formation

**DOI:** 10.1101/2022.01.12.475981

**Authors:** Leyao Shen, Yilin Yu, Yunji Zhou, Shondra M. Pruett-Miller, Guo-Fang Zhang, Courtney M. Karner

## Abstract

Cellular differentiation is associated with the acquisition of a unique protein signature which is essential to attain the ultimate cellular function and activity of the differentiated cell. This is predicted to result in unique biosynthetic demands that arise during differentiation. Using a bioinformatic approach, we discovered osteoblast differentiation is associated with increased demand for the amino acid proline. When compared to other differentiated cells, osteoblast-associated proteins including RUNX2, OSX, OCN and COL1A1 are significantly enriched in proline. Using a genetic and metabolomic approach, we demonstrate that the neutral amino acid transporter SLC38A2 acts cell autonomously to provide proline to facilitate the efficient synthesis of proline-rich osteoblast proteins. Genetic ablation of SLC38A2 in osteoblasts limits both osteoblast differentiation and bone formation in mice. Mechanistically, proline is primarily incorporated into nascent protein with little metabolism observed. Collectively, these data highlight a requirement for proline in fulfilling the unique biosynthetic requirements that arise during osteoblast differentiation and bone formation.

## Background

The mammalian boney skeleton is a remarkable organ that has multiple functions including support, mobility, protection of internal organs, endocrine signaling, mineral storage as well as being a site for red blood cell production (Guntur & Rosen, 2012; Jagannathan-Bogdan & Zon, 2013; Long, 2012; Salhotra, Shah, Levi, & Longaker, 2020). The skeleton develops embryonically through two distinct mechanisms, intramembranous and endochondral ossification (Berendsen & Olsen, 2015). Intramembranous ossification is responsible for forming the ‘flat’ bones of the skull. Here, mesenchymal progenitor cells condense and give rise to bone directly. The remainder of the skeleton develops through endochondral ossification. In this process, the mesenchymal progenitors condense and give rise to a cartilaginous template which is subsequently ossified. Regardless of the developmental mechanism, skeletal development depends upon osteoblasts. Osteoblasts are secretory cells responsible for producing and secreting the Collagen Type 1 (COL1A1) rich extracellular bone matrix. Osteoblast differentiation is tightly regulated by the transcription factors RUNX2 and OSX (encoded by *Sp7)* (Ducy, Zhang, Geoffroy, Ridall, & Karsenty, 1997; Nakashima et al., 2002; Otto et al., 1997; Takarada et al., 2016). Genetic studies in mice demonstrate RUNX2 is essential for commitment to the osteoblast lineage as well as the transcriptional regulation of osteoblast marker genes (e.g., *Spp1* and *Bglap*) (Komori et al., 1997; Meyer, Benkusky, Lee, & Pike, 2014; Otto et al., 1997; Wu et al., 2014). OSX functions downstream of RUNX2 to regulate osteoblast differentiation and osteoblast gene expression (e.g., *Spp1*, *Ibsp*, and *Bglap*) (Bianco, Fisher, Young, Termine, & Robey, 1991; Ducy et al., 1996).

During differentiation, osteoblasts acquire a distinct protein profile in addition to increasing bone matrix production (Alves et al., 2010; A. X. Zhang et al., 2007). Protein and bone matrix production is biosynthetically demanding and predicted to present differentiating osteoblasts with changing metabolic demands (Buttgereit & Brand, 1995). Thus, osteoblasts must maximize nutrient and amino acid acquisition for differentiation and matrix production to proceed. Consistent with this, both glucose and amino acid uptake are required for osteoblast differentiation and bone formation (Elefteriou et al., 2006; Rached et al., 2010; Wei et al., 2015). Osteoblasts primarily rely on glycolytic metabolism of glucose which provides ATP for protein synthesis and to regulate RUNX2 stability to promote osteoblast differentiation (Esen et al., 2013; W.-C. Lee, Ji, Nissim, & Long, 2020; Wei et al., 2015). Like glucose, amino acids have long been recognized as important regulators of osteoblast differentiation and bone matrix production (Elefteriou et al., 2006; Hahn, Downing, & Phang, 1971; Karner, Esen, Okunade, Patterson, & Long, 2015; Rached et al., 2010; Shen, Sharma, Yu, Long, & Karner, 2021; Yu et al., 2019). Affecting the ability of cells to sense or obtain amino acids either by limiting their availability in the media or inhibiting cellular uptake has detrimental effects on osteoblast differentiation and bone formation (Chen & Long, 2018; Elefteriou et al., 2006; Esen et al., 2013; Hu et al., 2020; Karner et al., 2015; Rached et al., 2010; Shen et al., 2021; Yu et al., 2019). Despite this, the role of individual amino acids in osteoblasts is not well understood. Recent studies identified glutamine as a particularly important amino acid in osteoblasts supporting protein and amino acid synthesis, redox regulation and energetics (Karner et al., 2015; Shen et al., 2021; Stegen et al., 2020; Yu et al., 2019). Whether other individual amino acids are similarly important for osteoblast differentiation remains unknown.

Proline is an intriguing amino acid in osteoblasts as it is important for both the biosynthesis and structure of collagen (Grant & Prockop, 1972; Krane, 2008). In addition, interest in proline has recently increased as proline is critical for cancer cell survival, tumorigenesis and metastasis (Liu, Glunde, et al., 2012; Nagano et al., 2017; Phang, Liu, Hancock, & Christian, 2012). Proline is a multifunctional amino acid with important roles in carbon and nitrogen metabolism, oxidative stress protection, cell signaling, nutrient adaptation and cell survival (Hollinshead et al., 2018; Liu, Le, et al., 2012; Phang, 2019). Proline can contribute to protein synthesis directly through incorporation into protein or can be metabolized into downstream products involved in energetic and biosynthetic reactions. Despite its emerging role in cancer cells, the role of proline during osteoblast differentiation and bone development is understudied.

Here we identify proline as a critical nutrient in osteoblasts. Using a multifaceted approach, we demonstrate that sodium-dependent neutral amino acid transporter-2 (SNAT2, encoded by *Slc38a2* and denoted herein as SLC38A2) acts cell autonomously to provide proline necessary for osteoblast differentiation and bone development. Mechanistically, proline is essential for the synthesis of proline-rich osteoblast proteins including those that regulate osteoblast differentiation (e.g., RUNX2 and OSX) and bone matrix production (e.g., COL1A1). These data highlight a broad requirement for proline to fulfill unique synthetic demands associated with osteoblast differentiation and bone formation.

## Results

### Proline is enriched in osteoblast-associated proteins, leading to increased proline demand during osteoblast differentiation

To identify if there are unique requirements for individual amino acids that arise during differentiation, we first profiled the amino acid composition of select proteins (e.g., RUNX2, OSX, COL1A1 and OCN) that are induced during osteoblast differentiation (Figure 1 Supplement 1). These classical osteoblast proteins are enriched with the amino acid proline and to a lesser extent alanine when compared to all proteins (Figure 1A and Table 1). For comparison, other amino acids were either uniformly underrepresented (e.g., Glu, Ile and Val) or were enriched only in a subset of these proteins (e.g., Gly and Gln) (Figure 1A and Table 1). To determine if this observation was characteristic of osteoblast proteins in general, we next evaluated amino acid enrichment in proteins that are associated with osteoblast differentiation based on gene ontology. Osteoblast-associated proteins (GO:0001649) were found to have a higher proline composition when compared to the average of all proteins (7.1% vs 6.1% proline for osteoblasts vs all proteins) (Table 2). In fact, many classical osteoblast proteins (e.g., RUNX2, OSX and COL1A1) were above the 90^th^ percentile for proline composition and 43.5% of all osteoblast proteins were above the 75^th^ percentile for proline composition. No other amino acids were similarly enriched in osteoblast-associated proteins (Table 2). Moreover, osteoblast-associated proteins were enriched for proline when compared to proteins associated with other cell types including osteoclasts (GO:0030855), cardiomyocytes (GO:0001649), muscle cells (GO:0055007), hematopoietic stem cells (GO:0042692), endothelial cells (GO:0030182), epithelial cells (GO:0055007) or neurons (GO:0030182) (Figure 1B and Table 2). In contrast, alanine enrichment was comparable amongst the different cell types (Figure 1B and Table 2). These data suggest that osteoblast differentiation is associated increased proline demand. To test this hypothesis, we transcriptionally profiled naïve calvarial cells that were induced to undergo osteoblast differentiation and quantified the proline enrichment of the encoded proteins. Consistent with our previous analysis, proline was enriched in proteins encoded by the induced genes compared to either all genes or genes that were suppressed in differentiated calvarial osteoblasts (Figure 1C). Moreover, comparing the basal and differentiation associated transcriptional changes with proline composition indicates that proline demand is predicted to rise whereas alanine demand is predicted to decline during osteoblast differentiation (Figure 1D). All together, these data predict proline is uniquely required during osteoblast differentiation due to the increasing expression of proline-enriched osteoblast proteins.

**Figure 1.**
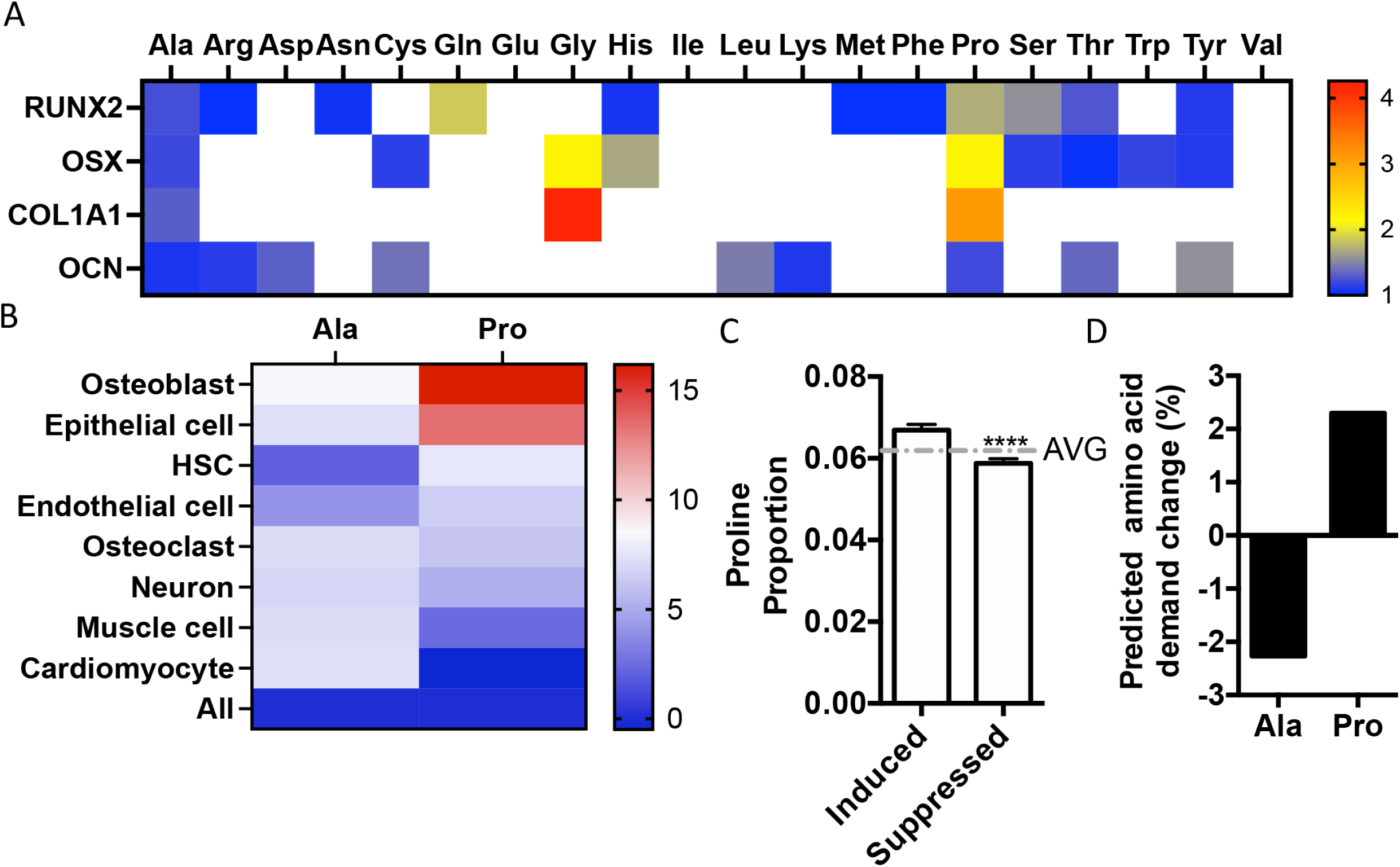
Osteoblast proteins are enriched with the amino acid proline. **(A)** Heat map depicting the relative amino acid enrichment for the indicated osteoblast proteins. Color bar represents fold enrichment relative to the average amino acid content. White boxes denote below average enrichment. **(B)** Heat map depicting alanine or proline enrichment in differentiation associated proteins. Color bar represents the percent increase in abundance relative to all proteins. **(C)** Graphical depiction of the proline proportion of the top 500 genes that are induced or suppressed during osteoblast differentiation. Dashed line represents the average proline proportion of all proteins. **** p≤ 0.00005. by unpaired 2-tailed Student’s t-test. **(D)** Graphical depiction of the predicted change in demand for alanine or proline based on changes in gene expression during osteoblast differentiation.

**Table 1.** Amino acid composition of classical osteoblast proteins.

|  | **RUNX2** | **OSX** | **COL1A1** | **OCN** | **ALL**<br>**PROTEINS** |
| --- | --- | --- | --- | --- | --- |
| **Ala** | 0.084 | 0.082 | 0.089 | 0.074 | 0.068 |
| **Cys** | 0.012 | 0.026 | 0.012 | 0.032 | 0.023 |
| **Asp** | 0.044 | 0.033 | 0.041 | 0.063 | 0.048 |
| **Glu** | 0.021 | 0.044 | 0.052 | 0.063 | 0.069 |
| **Phe** | 0.038 | 0.023 | 0.018 | 0.021 | 0.038 |
| **Gly** | 0.053 | 0.138 | 0.268 | 0.053 | 0.063 |
| **His** | 0.028 | 0.044 | 0.006 | 0.000 | 0.026 |
| **Ile** | 0.021 | 0.012 | 0.017 | 0.042 | 0.045 |
| **Lys** | 0.031 | 0.051 | 0.038 | 0.063 | 0.057 |
| **Leu** | 0.059 | 0.084 | 0.035 | 0.147 | 0.100 |
| **Met** | 0.023 | 0.012 | 0.010 | 0.021 | 0.023 |
| **Asn** | 0.038 | 0.026 | 0.023 | 0.032 | 0.036 |
| **Pro** | 0.105 | 0.133 | 0.190 | 0.074 | 0.061 |
| **Gln** | 0.089 | 0.033 | 0.033 | 0.032 | 0.048 |
| **Arg** | 0.056 | 0.040 | 0.047 | 0.063 | 0.056 |
| **Ser** | 0.133 | 0.098 | 0.046 | 0.074 | 0.085 |
| **Thr** | 0.069 | 0.056 | 0.030 | 0.074 | 0.054 |
| **Val** | 0.054 | 0.021 | 0.029 | 0.032 | 0.061 |
| **Trp** | 0.010 | 0.014 | 0.004 | 0.000 | 0.012 |
| **Tyr** | 0.030 | 0.030 | 0.010 | 0.042 | 0.027 |

**Table 2.** Relative amino acid composition of proteins associated with various differentiated cell types based on GO Terms.

|  | Osteoblast | Epithelial cell | HSC | Endothelial cell | Osteoclast | Neuron | Muscle cell | Cardiomyocyte | All |
| --- | --- | --- | --- | --- | --- | --- | --- | --- | --- |
| GO Term | 00001649 | 0030855 | 0030097 | 0045446 | 0030316 | 0030182 | 0042692 | 0055007 |  |
| **Ala** | 0.0739 | 0.0731 | 0.0695 | 0.0709 | 0.0730 | 0.0728 | 0.0730 | 0.0731 | 0.0681 |
| **Cys** | 0.0301 | 0.0230 | 0.0234 | 0.0259 | 0.0292 | 0.0228 | 0.0233 | 0.0223 | 0.0227 |
| **Asp** | 0.0478 | 0.0483 | 0.0493 | 0.0467 | 0.0468 | 0.0502 | 0.0511 | 0.0490 | 0.0479 |
| **Glu** | 0.0639 | 0.0678 | 0.0671 | 0.0682 | 0.0614 | 0.0693 | 0.0727 | 0.0690 | 0.0694 |
| **Phe** | 0.0331 | 0.0331 | 0.0365 | 0.0342 | 0.0374 | 0.0347 | 0.0367 | 0.0373 | 0.0375 |
| **Gly** | 0.0686 | 0.0698 | 0.0672 | 0.0685 | 0.0656 | 0.0667 | 0.0666 | 0.0667 | 0.0629 |
| **His** | 0.0273 | 0.0256 | 0.0266 | 0.0239 | 0.0245 | 0.0254 | 0.0249 | 0.0256 | 0.0262 |
| **Ile** | 0.0362 | 0.0401 | 0.0409 | 0.0458 | 0.0407 | 0.0420 | 0.0429 | 0.0441 | 0.0445 |
| **Lys** | 0.0531 | 0.0557 | 0.0570 | 0.0531 | 0.0499 | 0.0560 | 0.0606 | 0.0627 | 0.0571 |
| **Leu** | 0.0946 | 0.0923 | 0.0957 | 0.0950 | 0.1040 | 0.0961 | 0.0930 | 0.0903 | 0.1004 |
| **Met** | 0.0220 | 0.0232 | 0.0229 | 0.0224 | 0.0210 | 0.0222 | 0.0222 | 0.0238 | 0.0228 |
| **Asn** | 0.0360 | 0.0361 | 0.0362 | 0.0382 | 0.0366 | 0.0374 | 0.0369 | 0.0363 | 0.0360 |
| **Pro** | 0.0711 | 0.0694 | 0.0659 | 0.0652 | 0.0650 | 0.0644 | 0.0627 | 0.0609 | 0.0612 |
| **Gln** | 0.0469 | 0.0472 | 0.0468 | 0.0458 | 0.0446 | 0.0459 | 0.0464 | 0.0482 | 0.0478 |
| **Arg** | 0.0607 | 0.0573 | 0.0563 | 0.0566 | 0.0539 | 0.0575 | 0.0561 | 0.0547 | 0.0559 |
| **Ser** | 0.0853 | 0.0874 | 0.0848 | 0.0807 | 0.0843 | 0.0835 | 0.0797 | 0.0848 | 0.0853 |
| **Thr** | 0.0527 | 0.0529 | 0.0537 | 0.0557 | 0.0563 | 0.0538 | 0.0528 | 0.0544 | 0.0543 |
| **Val** | 0.0568 | 0.0588 | 0.0599 | 0.0639 | 0.0623 | 0.0598 | 0.0599 | 0.0585 | 0.0610 |
| **Trp** | 0.0119 | 0.0110 | 0.0122 | 0.0119 | 0.0135 | 0.0117 | 0.0108 | 0.0105 | 0.0119 |
| **Tyr** | 0.0278 | 0.0279 | 0.0280 | 0.0275 | 0.0298 | 0.0279 | 0.0275 | 0.0278 | 0.0270 |

We next sought to understand proline dynamics in osteoblasts. Proline can be taken up from the extracellular milieu or synthesized. To determine the source of proline in osteoblasts, we first performed stable isotopomer analysis using ^13^C_U_-Proline to evaluate proline uptake or either ^13^C_U_-Glutamine or ^13^C_1,2_-Glucose to estimate *de novo* proline biosynthesis. 10.5% of intracellular proline is synthesized from either glutamine (9.9%) or glucose (0.6%) in 24 hours (Figure 2A). By comparison, 37.8% of the proline pool is labeled from ^13^C_U_-proline after 24 hours and this increased to 66.6% after 72 hours (Figure 2A). The slow labeling of proline when compared to intracellular glutamine which reached saturation within hours, suggests that in naïve calvarial cells, proline uptake is slow, and the intracellular proline pool is relatively stable with little turnover. To test this, we performed radiolabeled amino acid uptake assays to compare the rates of proline and glutamine uptake. Consistent with the labeling data, proline uptake was slow compared to glutamine uptake in naïve cells (Figure 2B and Figure 2 Supplement 1A). During differentiation, the rate of proline uptake increased significantly and to a greater extent than glutamine which also increased (Figure 2C and Figure 2 Supplement 1B). The tracing experiments indicated little proline metabolism occurs in osteoblasts as proline carbon was not observed in any other amino acid or downstream metabolite even after 72 hours (Figure 2A). By comparison, carbon from both glutamine and glucose was observed in many metabolites including TCA cycle intermediates and amino acids (Figure 2A and not shown). These data suggest proline is not metabolized and is primarily used for protein synthesis. Consistent with this conclusion, proline incorporation into both total protein and collagen significantly increases during differentiation (Figure 2D). Moreover, almost 50% of proline in total protein was derived from ^13^C_U_-Proline (Figure 2E). Importantly, we observed no proline-derived amino acids in total protein despite the presence of glutamine derived amino acids including proline (Figure 2E). Thus, proline demand and protein synthesis rise concomitantly during osteoblast differentiation.

**Figure 2.**
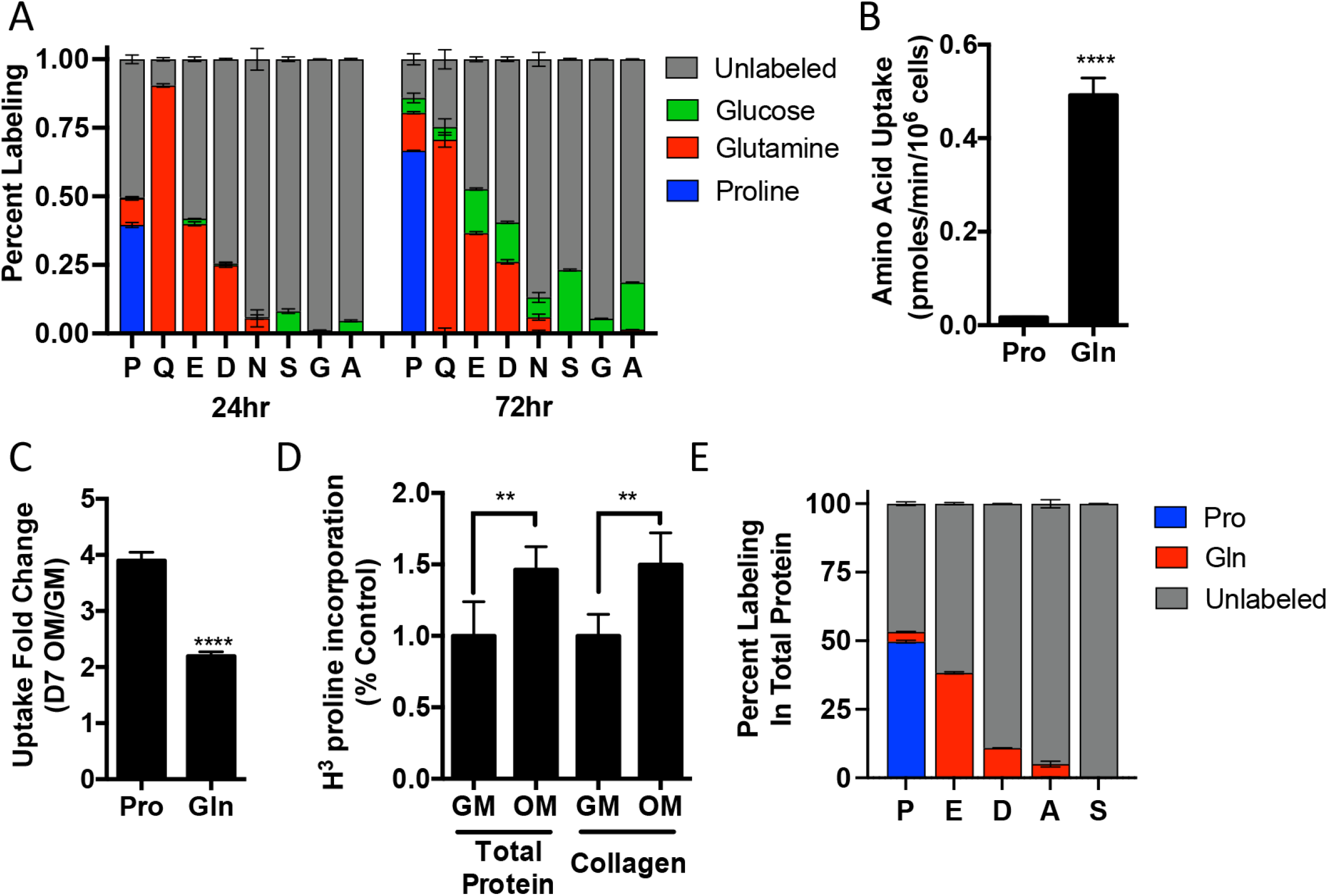
Proline uptake and incorporation into protein increases during osteoblast differentiation. **(A)** Graphical depiction of proline, glutamine, glutamate, aspartate, asparagine, serine, glycine and alanine labeling from [U-^13^C]-Proline (n=3), [U-^13^C]-Glutamine (n=3) or [1,2-^13^C]-Glucose (n=3) in naïve calvarial osteoblasts. **(B-C)** Radiolabeled ^3^H-Proline uptake assay performed in naive bone marrow stromal cells (BMSC) **(B)** or after 7 days of osteoblast differentiation **(C)**. **(D)** Contribution of [U-^13^C]-Proline or [U-^13^C]-Glutamine to proline, glutamate, aspartate, alanine or serine isolated from total protein (n=3). **(E)** Radiolabeled proline incorporation assay performed in primary calvarial cells cultured in growth medium (GM) or osteogenic medium (OM) for 7 days (n=3). ** p≤ 0.005, **** p≤ 0.00005. by unpaired 2-tailed Student’s t-test.

We next sought to determine the effects of proline withdrawal on protein expression. Proline withdrawal specifically reduced charging of the proline tRNA (AGG) but did not affect the activation of either the mTOR pathway (as determined by S6 ribosomal protein phosphorylation at S235/236) or the integrated stress response (ISR) (as determined by and EIF2a phosphorylation at Ser51) (Figure 3 Supplement 1A-B). Proline withdrawal did not affect the expression of select non-proline enriched proteins (Figure 3A and Figure 3 Supplement 1B). Conversely, proline withdrawal significantly reduced the expression of osteoblast proteins that had higher than average proline content including COL1A1 (19.1% proline), RUNX2 (10.5% proline), OSX (13.3% proline) and ATF4 (10.6% Proline) (Figure 3A). Importantly, proline withdrawal did not affect the mRNA expression of these proteins (Figure 3 Supplement 1C). We next took a candidate approach and evaluated other proline-enriched (e.g., EIF4EBP1 (13.7% proline), PAX1 (11.1% proline), ATF2 (10.7% proline), SMAD1 (9.9% proline), and EIF2A, (7.6% proline)) and non-enriched proteins (ERK1 (6.6% proline), PHGDH (5.3% proline), EEF2 (5.2% proline), AKT (4.6% proline), ACTB (5.1% proline), mTOR (4.4% proline), TUBA (4.4% proline) and S6RP (4.2% proline)) that are not known to be required for osteoblast differentiation but are expressed in calvarial cells according to our transcriptomic analyses. Proline withdrawal significantly reduced the expression of the proline enriched proteins without affecting the low proline proteins (Figure 3A and Figure 3 Supplement 1B). The reduction in protein expression significantly correlated with the proline content in the proteins (Figure 3B). These data indicate this phenomenon is broadly generalizable in osteoblasts. The decreased protein expression is due primarily to reduced synthesis of proline enriched proteins as cycloheximide washout experiments found proline withdrawal resulted in a significant delay in the recovery of proline-enriched protein expression (Figure 3C and Figure 3 Supplement 1D-H). Thus, proline is essential for the synthesis of proline-enriched osteoblast proteins.

**Figure 3.**
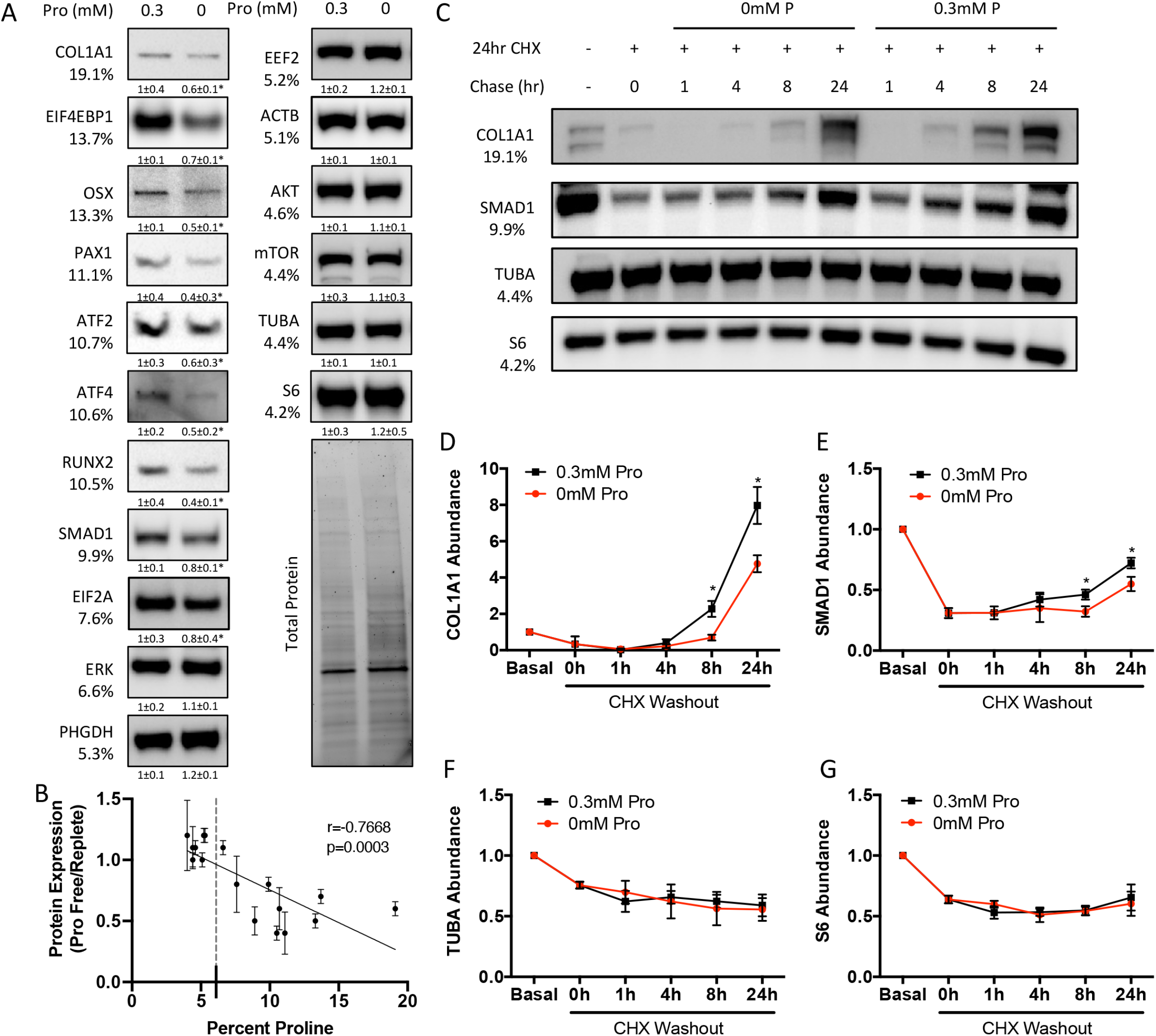
Proline is essential for the synthesis of proline-enriched osteoblast proteins. **(A)** Western blot analysis of naïve calvarial cells cultured in 0.3mM or 0mM Pro for 48 hours (n=3). In all blots, the percent proline composition is noted under the protein name. Protein expression normalized to total protein. Fold change ± SD for three independent experiments. **(B)** Correlation analysis of protein expression as a function of the proline composition of proteins in naïve calvarial cells cultured in media containing either 0mM or 0.3mM Pro for 48 hours. **(C-G)** The effect of proline availability on the synthesis of select proteins. CHX – cycloheximide. Error bars depict SD. * p≤ 0.05. by unpaired 2-tailed Student’s t-test.

### SLC38A2 provides proline to facilitate the synthesis of proline rich osteoblast proteins

We next sought to identify the proline transporter in osteoblasts. Proline uptake in osteoblasts is reported to occur in a 2-(methylamino)-isobutyric acid (MeAIB) sensitive manner (Baum & Shteyer, 1987; Yee, 1988). Consistent with these reports, MeAIB reduced proline uptake by 80% in both osteoblasts and bone shafts with minimal effects on the uptake of other amino acids (e.g., Gln, Ala, Gly or Ser) (Figure 4A and Figure 4 Supplement 1A). We next sought to identify candidate proline transporters based on relative mRNA expression. Evaluation of our transcriptomic data identified *Slc38a2* as the highest expressed putative proline transporter in calvarial cells (Table 3). *Slc38a2* encodes the sodium-dependent neutral amino acid transporter-2 (SNAT2, denoted here as SLC38A2) which transports neutral alpha amino acids (e.g., proline) in a Na^+^ dependent manner that is sensitive to MeAIB (Grewal et al., 2009; Hoffmann et al., 2018). To determine if SLC38A2 transports proline in differentiating osteoblasts, we targeted *Slc38a2* using a CRISPR/Cas9 approach (Figure 4 Supplement 1B). *Slc38a2* targeting significantly reduced SLC38A2 protein and reduced radiolabeled proline uptake by ∼50% in differentiated calvarial cells (Figure 4B-C). This is likely a slight underestimation of SLC38A2 dependent proline uptake due to incomplete ablation of SLC38A2 protein (Figure 4C). Consistent with decreased proline uptake, *Slc38a2* ablation specifically reduced proline-tRNA charging similar to proline withdrawal without negatively affecting charging of other tRNAs or activating the ISR (Figure 4 Supplement 1C-D). Moreover, *Slc38a2* ablation specifically reduced the expression of the proline enriched proteins without affecting the expression of non-proline enriched proteins or mRNA expression of these proteins (Figure 4C and Figure 4 Supplement 1D). The effect of *Slc38a2* ablation on protein expression significantly correlated with the proline content in the proteins (Figure 4D). This is likely a direct result of decreased proline uptake as *Slc38a2* deletion did not affect mTOR activation or induce ISR (Figure 4 Supplement 1D). Thus, SLC38A2 provides proline for the efficient synthesis of proline-enriched proteins.

**Table 3.** mRNA expression of putative proline transporters.

|  | **System** | **Alias** | **cOB**<br>**FPKM** |
| --- | --- | --- | --- |
| **Slc38a2** | A | SNAT2 | 8823.1 |
| **Slc1a4** | ASC | ASCT1 | 3030.7 |
| **Slc36a4** | LYAAT | PAT4 | 1929 |
| **Slc36a1** | LYAAT | PAT1 | 803.7 |
| **Slc38a4** | A | SNAT4 | 267.9 |
| **Slc36a2** | LYAAT | PAT2 | 1.1 |
| **Slc6a15** | B<sup>0</sup> | B<sup>0</sup>AT2 | 7.5 |
| **Slc36a3** | LYAAT | PAT3 | 0 |
| **Slc6a7** | IMINO<sup>B</sup> | PROT | 4.3 |
| **Slc6a20a** | IMINO | SIT2 | 0 |
| **Slc6a20b** | IMINO | SIT1 | 0 |
| **Slc6a19** | B<sup>0</sup> | B<sup>0</sup>AT1 | 0 |

**Figure 4.**
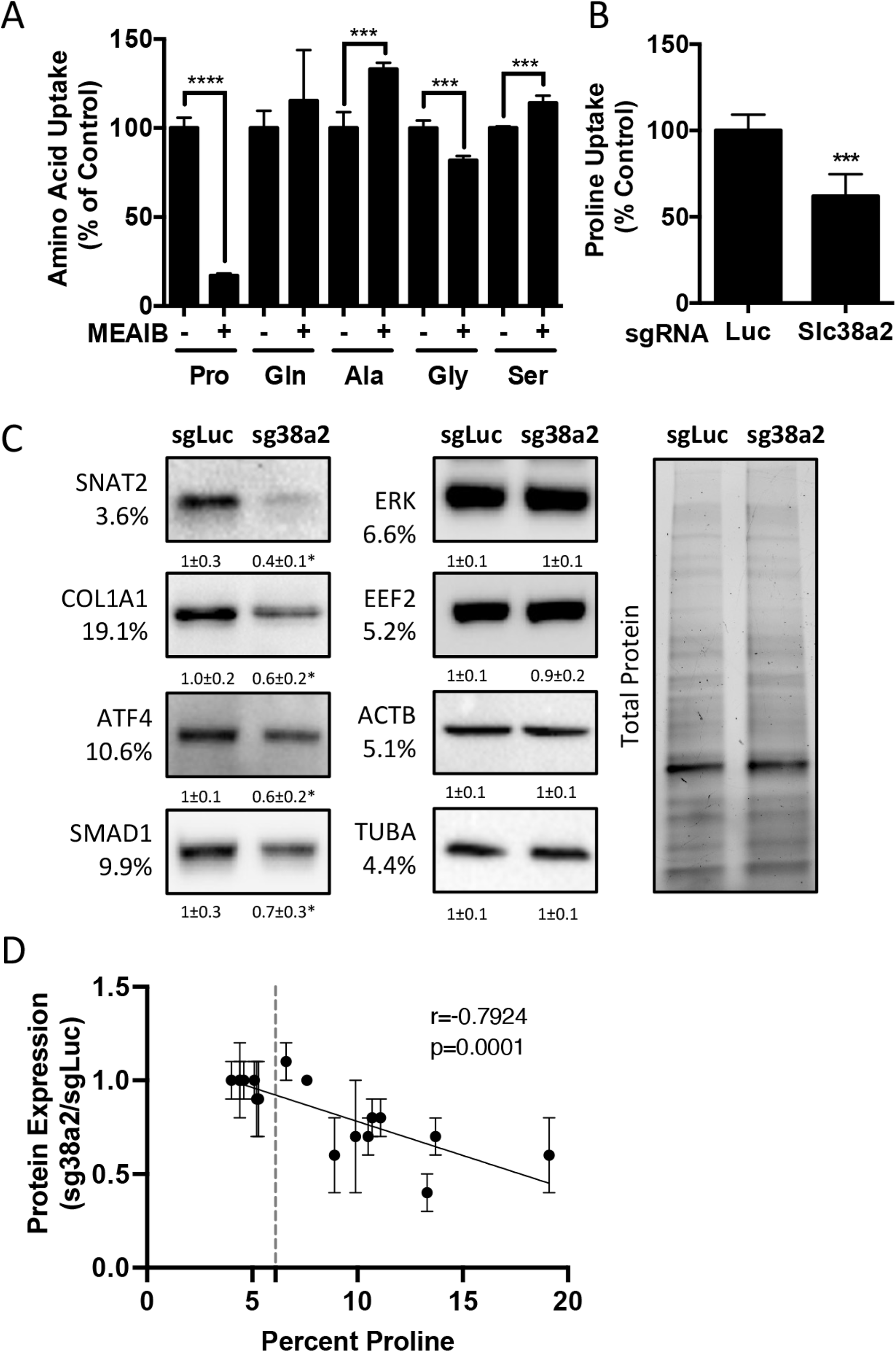
*Slc38a2* provides proline critical for the synthesis of proline-rich proteins. **(A)** Graphical depiction of the effects of 5mM MEAIB on radiolabeled amino acid uptake in primary calvarial cells (n=3). **(B,C)** Effect of *Slc38a2* targeting on ^3^H-Proline uptake (n=5) **(B)**, or protein expression **(C)**. In all blots, the percent proline composition is noted under the protein name Protein expression normalized to total protein. Fold change ± SD for three independent experiments. **(D)** Correlation analysis of protein expression as a function of the proline composition of proteins in wild type (sgLuc) or *Slc38a2* targeted calvarial cells (sg38a2) calvarial cells. * p≤ 0.05, *** p≤ 0.0005, **** p≤ 0.00005. by unpaired 2-tailed Student’s t-test.

### *SLC38A2* provides proline necessary for bone development

We next sought to understand the role of SLC38A2 during osteoblast differentiation. *Slc38a2* deletion did not affect cell viability, proliferation, or the mRNA expression or induction of early osteoblast regulatory genes (e.g., *Akp2* and *Runx2*) (Figure 5 Supplement 1A-C). However, *Slc38a2* deficient cells were characterized by reduced induction of *Sp7* and terminal osteoblast marker genes (e.g., *Ibsp* and *Bglap*) as well as reduced matrix mineralization (Figure 5 Supplement 1C-D). This indicates that SLC38A2 provides proline required for terminal osteoblast differentiation and matrix mineralization *in vitro*.

In light of these data, we next analyzed the function of *Slc38a2* during osteoblast differentiation by comparing mice null for SLC38A2 due to the insertion of LacZ into the coding region of *Slc38a2* (*Slc38a2^LacZ/LacZ^*). We verified the absence of SLC38A2 expression by western blot (Figure 5 Supplement 2A). Using alcian blue and alizarin red staining (which stains cartilage and bone matrix blue or red respectively) we found that *Slc38a2^LacZ/LacZ^* embryos were characterized by a conspicuous reduction in red mineralized bone matrix staining in both endochondral and intramembranous bones at E15.5 (Figure 5A-B). This defect in bone mineralization was most obvious in the developing skull (Figure 5B). By comparison, Slc38a2^LacZ/LacZ^ animals had no apparent defects in cartilage formation at E15.5 indicating loss of *Slc38a2* impacts osteoblast differentiation. To test this, we crossed mice harboring a floxed allele of *Slc38a2* (*Slc38a2^fl^*) with mice expressing Cre recombinase under the control of the *Sp7* promoter (*Sp7Cre*) which is active in osteoblast progenitors beginning at E14.5 (Rodda & McMahon, 2006). *Sp7Cre;Slc38a2^fl/fl^* bones were characterized by reduced SLC38A2 expression and reduced proline uptake (Figure 5 Supplement 3A-B). Like the *Slc38a2^LacZ/LacZ^* mice, *Sp7Cre;Slc38a2^fl/fl^* mice had significantly less alizarin red stained bone matrix at E15.5 (Figure 5C-D). By postnatal day 1 (P1), overall bone matrix in long bones was comparable in both genetic models, however the skulls from both *Slc38a2^LacZ/LacZ^* and *Sp7Cre;Slc38a2^fl/fl^* mice continued to be poorly mineralized with patent fontanelles compared to their respective littermate controls (Figure 5E-H, Figure 5 Supplement 3C). Because we observed a consistent defect in intramembranous ossification and *Sp7Cre* is expressed in both osteoblasts and hypertrophic chondrocytes in the developing limbs we focused our molecular analyses on the osteoblasts in the developing calvarium. *Sp7Cre;Slc38a2^fl/fl^* calvariae had normal alkaline phosphatase staining despite less mineralized area shown by von Kossa staining (Figure 5I-L). The defects in bone development are attributed to delayed osteoblast differentiation as *Sp7Cre;Slc38a2^fl/fl^* mice had significantly reduced expression of *Spp1*, *Ibsp* and *Bglap* (Figure 5M-R). This was not due to a reduction in overall osteoblast numbers as there was no difference in the total number of *Sp7:GFP* expressing cells per mineralized area (Figure 5S-T). Despite this, significantly fewer *Sp7:*GFP expressing cells were found to have OSX (encoded by *Sp7*) or RUNX2 protein expression in *Sp7Cre;Slc38a2^fl/fl^* animals (Figure 5S-T and Figure 5 Supplement 3C). Similar results were observed in the limbs of both *Slc38a2^LacZ/LacZ^* and *Sp7Cre;Slc38a2^fl/fl^* mice at E15.5 (Figure 5 Supplements 2B and 3D). Similarly, *Sp7GFP* expressing cells in *Sp7Cre;Slc38a2^fl/fl^* mice had significantly reduced COL1A1 protein expression despite normal *Col1a1* mRNA expression when compared to *Sp7Cre;Slc38a2^fl/+^* controls (Figure 5U-X and Figure 5 Supplement 2B and 3C). For comparison, the expression of proline poor Actin (as determined by phalloidin staining) and GFP (4.2% Proline) were unaffected in *Sp7Cre;Slc38a2^fl/fl^* calvariae (Figure 5 Supplement 3C). Collectively these data indicate *Slc38a2* provides proline essential for osteoblast differentiation and bone formation during bone development.

**Figure 5.**
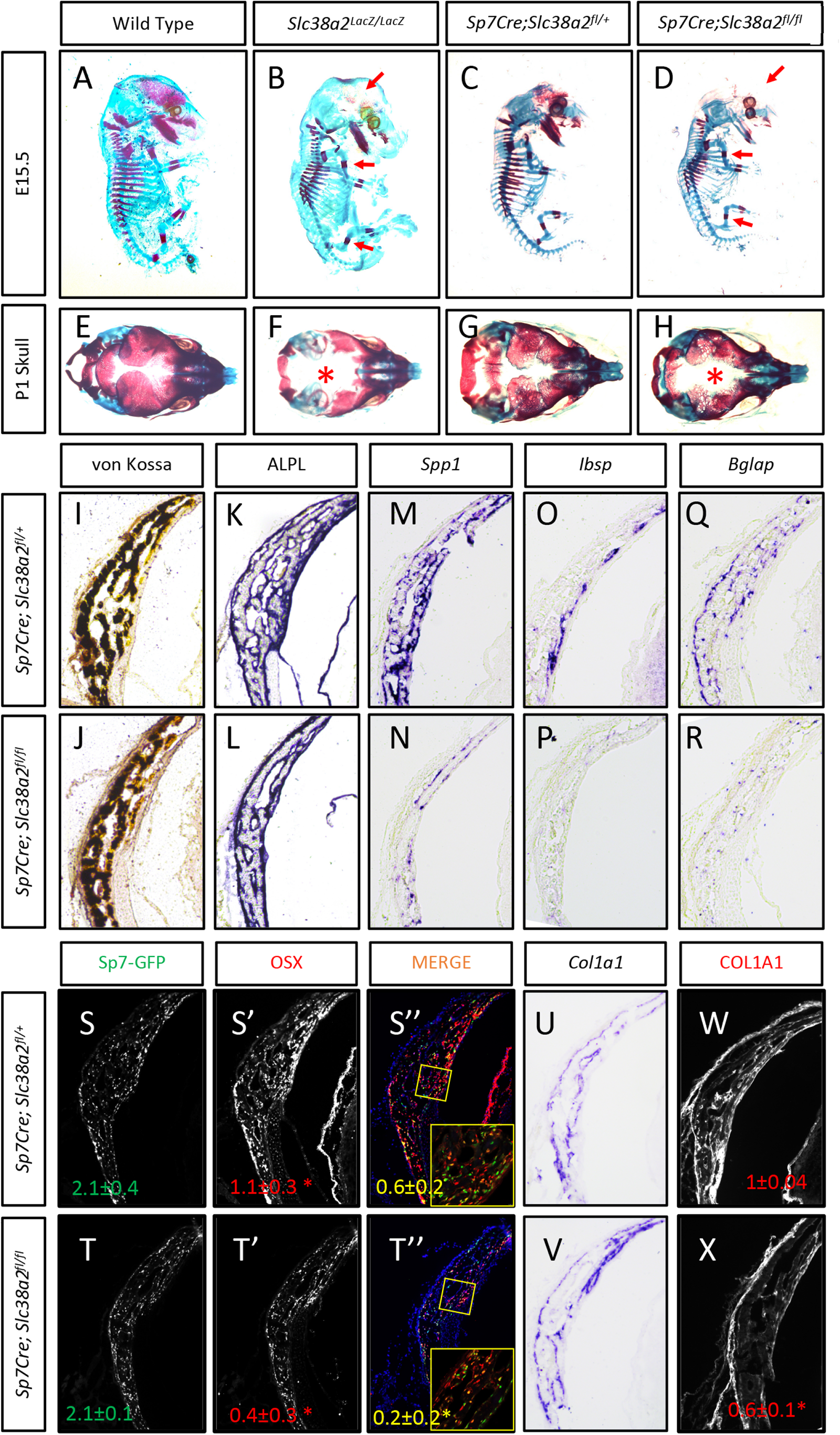
*Slc38a2* dependent proline uptake is required for osteoblast differentiation during bone development. **(A-H)** Skeletal preparations of *Slc38a2^LacZ/LacZ^* or wildtype controls **(A-B, E-F)** or *Sp7Cre;Slc38a2^fl/fl^* or *Sp7Cre;Slc38a2^fl/+^* littermate controls **(C-D, G-H)** at E15.5 **(A-D)** or P1 **(E-H)**. Red arrow **(A-D)** or Asterix **(E-H)** highlights reduced mineralization. A total of n=7 or n=5 *Slc38a2^LacZ/LacZ^* animals and n=5 or n=5 for *Sp7Cre;Slc38a2^fl/fl^* animals were analyzed at E15.5 or P1 respectively. **(I-R)** Representative von Kossa staining **(I,J)**, alkaline phosphatase (ALPL) staining **(K,L**) *in situ* hybridization for *Spp1* **(M,N)***, Ibsp* **(O,P)**, *Bglap* **(Q,R)** and *Col1a1* **(U,V)**, or immunofluorescent staining for OSX **(S’,S”,T’,T”)** and COL1A1 **(W,X)** on *Sp7Cre;Slc38a2^fl/fl^* **(J,L,N,P,R,T,V,X)** or *Sp7Cre;Slc38a2^fl/+^* **(I,K,M,O,Q,S,U,W)** newborn calvariae. *p≤0.05. by paired 2-tailed Student’s t-test.

## Discussion

Here we have defined the major role for proline during osteoblast differentiation and bone formation. Namely, that osteoblasts require proline to fulfill unique biosynthetic demands that arise due to increased production of proline enriched osteoblast-associated proteins. Consistent with this, osteoblasts significantly increase proline consumption and to a lesser extent proline biosynthesis during differentiation. Genetically limiting proline uptake by ablating the proline transporter SLC38A2 results in delayed bone development and decreased bone mass in adult male mice. Mechanistically, osteoblasts utilize proline primarily for the synthesis of proline enriched osteoblast proteins to facilitate both osteoblast differentiation and bone matrix production. Collectively, these data highlight a broad requirement for proline to regulate osteoblast differentiation and bone development in addition to supporting collagen synthesis.

Osteoblast differentiation is characterized by a distinct protein profile in addition to increasing bone matrix production (Alves et al., 2010; A. X. Zhang et al., 2007). These osteoblast-associated proteins are enriched for the amino acid proline compared to all other proteins (Figure 1 and Tables 1-2). We and others have recently described increased consumption of numerous amino acids in differentiating osteoblasts including glutamine (Sharma, Yu, Shen, Zhang, & Karner, 2021; Stegen et al., 2020; Yu et al., 2019), asparagine (Sharma et al., 2021) and proline (this study). Glutamine and asparagine contribute to both *de novo* amino acid biosynthesis and protein synthesis directly (Sharma et al., 2021). Our data here indicates the primary use for proline is direct incorporation into nascent protein. Consistent with this, reducing proline availability specifically reduced the synthesis of proteins with higher-than-average proline content without affecting mTOR activation or inducing ISR (Figures 3 and 4). In fact, protein expression was negatively correlated with the proline content in proteins when proline availability or uptake was limited (Figure 3 and 4). By comparison, limiting the availability of glutamine induced robust activation of the ISR and inhibits global protein synthesis (Sharma et al., 2021). This likely reflects the necessity of glutamine metabolism to maintain amino acid concentrations (including proline) and provide other metabolites during osteoblast differentiation (Sharma et al., 2021; Stegen et al., 2020; Yu et al., 2019). Consistent with the more direct use of proline in protein but not amino acid biosynthesis, we did not observe activation of the ISR in proline free conditions despite reduced charging of proline tRNA. It is important to note that we evaluated the effects of proline withdrawal for 48 hours. This time point may miss the chronic effects of proline withdrawal as proline uptake is slow and the intracellular proline pool is stable with low turnover in naïve calvarial cells (Figure 2). Under these conditions, *de novo* biosynthesis of proline may be sufficient to meet the basal needs of naïve calvarial cells. Regardless, proline removal results in reduced synthetic efficiency of proline-rich proteins. This effect is likely exacerbated during osteoblast differentiation as these proline-rich proteins are increased.

Previous studies characterized proline uptake in both bones and osteoblasts directly. These studies described proline uptake occurring primarily via System A but did not identify individual transporters mediating proline uptake (Adamson & Ingbar, 1967; Finerman & Rosenberg, 1966; Hahn, Downing, & Phang, 1969; Yee, 1988). Here, we identified the sodium-dependent neutral amino acid transporter SLC38A2 as responsible for approximately 55% of proline uptake in both calvarial osteoblasts and isolated bones (Figure 4B and Figure 5 Supplement 3B). This is consistent with previous reports that System A mediates 60% of proline uptake in osteoblasts (Yee, 1988). Interestingly, SLC38A2 ablation affected only proline uptake (Figure 5 Supplement 3B). It is not clear why SLC38A2 exclusively transports proline in osteoblasts as amino acid transporters are thought to be promiscuous in their substrate specificity (Kandasamy, Gyimesi, Kanai, & Hediger, 2018; Teichmann et al., 2017). For example, SLC38A2 is reported to transport alanine, serine, glycine and glutamine in different cellular contexts (Bröer, Rahimi, & Bröer, 2016; Morotti et al., 2019). Our data indicates glutamine is not a primary substrate for SLC38A2 in bone cells. This is consistent with our recent data demonstrating glutamine uptake is mediated primarily by System ASC with no involvement of System A in osteoblasts (Sharma et al., 2021; Shen et al., 2021). In light of these data, a better understanding of the molecular regulation of SLC38A2 activity and substrate specificity is needed. In addition, it will be important to identify the transporters mediating SLC38A2 independent proline uptake as well as to understand their function during osteoblast differentiation and bone development.

Reducing proline uptake inhibited bone development in mice (Figure 5). This phenotype was attributed primarily to decreased osteoblast differentiation and reduced bone matrix production. Osteoblast differentiation and bone matrix production are associated with a unique biosynthetic demand for proline. Using a bioinformatic approach, we discovered that osteoblast associated proteins are more enriched for proline than any other amino acid when compared to other cell types, (Figure 1). Many of these proline-rich proteins are essential regulators of osteoblast differentiation (e.g. RUNX2, OSX and ATF4), bone matrix production (e.g., COL1A1) or regulate the endocrine functions of bone (e.g., OCN) (Ducy et al., 1996; Ducy et al., 1997; Elefteriou et al., 2006; Kern, Shen, Starbuck, & Karsenty, 2001; Nakashima et al., 2002; Otto et al., 1997; Yang et al., 2004). Limiting proline availability by genetically ablating SLC38A2 specifically affected the production of proline rich proteins (e.g., RUNX2, OSX and COL1A1) in a manner that was proportional to the relative proline content. It is important to note that relatively minor reductions in protein expression or function are known to negatively impact osteoblast differentiation and bone development and underly human bone diseases (Baek et al., 2013; Bardai et al., 2016; Ben Amor, Roughley, Glorieux, & Rauch, 2013; Choi et al., 2001; Lapunzina et al., 2010; B. Lee et al., 1997; Lou et al., 2009; Mundlos et al., 1997; S. Zhang et al., 2009). Thus, ablating SLC38A2 dependent proline uptake has broad effects on osteoblast differentiation due to minor reductions in many proline-rich osteoblast regulatory proteins. This highlights an unappreciated mechanism by which osteoblast differentiation is responsive to nutrient (e.g., proline) availability. When proline is available, osteoblast progenitors efficiently synthesize the proline rich proteins necessary for differentiation (RUNX2 and OSX) and bone matrix deposition (COL1A1). The high proline content of these proteins presents a novel cellular checkpoint to ascertain if appropriate resources, in this case proline, are available for osteoblast differentiation to proceed. When proline is limited, these proteins are not efficiently synthesized which limits osteoblast differentiation and bone matrix production until sufficient proline is available. This is critical to ensure cells can meet the synthetic challenges associated with osteoblast differentiation and bone matrix production.

In summary, we have defined the necessity and the molecular substrates of *Slc38a2* in osteoblasts. Our data indicates that SLC38A2 acts cell autonomously in osteoblasts to provide proline and that SLC38A2 is the major proline transporter in osteoblasts. Proline is essential for the production of proline-rich transcription factors (e.g., RUNX2 and OSX) and matrix proteins (COL1A1) necessary for osteoblast differentiation and bone formation. These data expand our understanding of the regulation of proline uptake and usage in osteoblasts and underscore the necessity of proline for osteoblast differentiation and bone development.

## Materials and Methods

### Mouse strains

*C57Bl/6J* (RRID: IMSR_JAX:000664), *Rosa26Cas9* (RRID: IMSR_JAX:024858), *Rosa26FLP* (RRID: IMSR_JAX:003946) and *Sp7-tTA,tetO-EGFP/Cre* (RRID: IMSR_JAX:006361) mouse strains were obtained from the Jackson Laboratory. *Slc38a2LacZ* (C57BL/6N-A<tmlBrd>Slc38a2<tm1a(KOMP)Wtsi>/Wtsi Ph) was purchased from the European Mouse Mutant Archive (www.emmanet.org). To generate *Slc38a2^flox^*, *Slc38a2^LacZ^* mice were crossed to *Rosa26FLP* to remove FRT-flanking LacZ cassette followed by a backcrossing with *C57Bl/6J* to remove *Rosa26^FLP^* allele. Mice were housed at 23 °C on a 12-hour light/dark cycle with free access to water and PicoLab Rodent Diet 20 (LabDiet #5053, St. Louis, MO). All mouse procedures were approved by the Animal Studies Committees at Duke University first and then the University of Texas Southwestern Medical Center at Dallas.

### Mouse analyses

Skeletal preparations were performed on embryonic day (E) 15.5 or postnatal (P) day 0 embryos obtained from timed pregnancies. Noon of the day of plugging was considered 12 hours post coitum or E0.5. Embryos are dehydrated in 95% ethanol overnight followed by submersion in acetone overnight. Specimens were then stained with 0.03% alcian blue in 70% ethanol and 0.005% alizarin red in water overnight. Stained embryos were then cleared in 1% KOH prior to a graded glycerol series (30%, 50% and 80%). For histological analyses, freshly isolated limbs or calvariae were fixed in 4% PFA at 4°C overnight. Limbs were then processed and embedded in paraffin and sectioned at 5*μ*m using a Microtome (Leica RM2255). Calvariae were cryoprotected in 30% sucrose overnight, embedded in OCT and sectioned at 10*μ*m using a Cryostat (Leica CM1950).

### *In Situ* hybridization

*In situ* hybridization was performed on 10µm cryosectioned calvariae or 5µm paraffin-sectioned limbs. Cryosections were washed with water first for 5min. Paraffin sections were deparaffinized and rehydrated, followed by 20mg/ml proteinase K treatment for 10min. Sections were first then fixed in 4%PFA for 10min followed by 10 min acetylation. Sections were then incubated in hybridization buffer for 2 hours at room temperature. Digoxigenin-labeled antisense RNA probes for *Col1a1* (HindIII, T7), *Sp7* (NotI, T3), *Spp1* (EcoRI, SP6), *Ibsp* (NOTI, SP6) or *Bglap* (XbaI, T3) were hybridized at 60 °C overnight.

### Immunohistochemistry

Sections were blocked in 1.5% goat serum in PBST and incubated with the following primary antibodies (1:250 in blocking solution) as indicated: Anti-Col1a1 (AB_1672342), Anti-Osx (AB_2194492) or Anti-Runx2 (AB_10949892) at 4 °C overnight. Sections were then incubated with Alexa Fluor 568 goat anti-rabbit (AB_143157)/-mouse IgG(H+L) antibody (AB_2534072) at 1:250 dilution at room temperature for 30min. Sections were post fixed in 4% PFA for 10 min before mounting. For actin staining, Alexa Fluor 647 Phalloidin (Invitrogen; 1:200 in blocking buffer) was applied to sections before mounting. Sections were mounted using Heatshield with DAPI (Vector).

### Cell culture

Primary calvarial osteoblasts were isolated as follows. The calvaria of P4 pups were harvested and extemporaneous tissue was removed. The calvariae was chopped by scissor into small pieces and washed with PBS twice. The calvaria pieces were then incubated in 1.8mg/mL Collagenase P in PBS for 10 minutes with agitation at 37°C four times. The first digestion was discarded, and the last three digestions were collected and run through 70 *μ*m cell strainer. Cells were then centrifuged at 350x g for 5min and cultured at T75 flasks in aMEM containing 15% FBS at 37 °C and 5% CO_2_. Cells were plated at 1x10^5^ cells/mL for further experiments when it reached 90% confluency. Osteoblast differentiation was induced at 100% confluency using aMEM supplemented with 50 mg/ml ascorbic acid and 10 mM β-glycerophosphate for the indicated time with a change of media every 48 hours. For proline drop out experiments, primary calvarial cells were treated with proline-free aMEM (Genaxxon) supplemented back to 0.3mM proline or not for the indicated length of time. To evaluate the synthesis of individual proteins, cycloheximide (CHX) washout experiments were performed. Calvarial osteoblasts were treated with 10*μ*g/mL CHX for 24 hours. Cells were then chased with aMEM containing either 0.3mM or 0mM proline for up to 24h before proteins were harvested. Alkaline phosphatase activity was assessed using 5-bromo-4-chloro-3’-indolyphosphate/nitro blue tetrazolium (BCIP/NPT). Mineralization was visualized by either von Kossa or Alizarin Red staining as indicated.

### CRISPR/Cas9 targeting

Lentiviral vectors expressing single guide RNAs (sgRNA) targeting either *Slc38a2* or Luciferase and mCherry were cloned into the LentiGuide-Puro plasmid according to the previously published protocol (Sanjana, Shalem, & Zhang, 2014). The LentiGuide-Puro plasmid was a gift from Feng Zhang (Addgene plasmid #52963). Sequences of each sgRNA protospacer are shown in Table S1. To make viral particles, the sgRNA carrying lentiviral vector was cotransfected in 293T cells with the plasmids pMD2.g and psPax2. Virus containing media was collected and run through 0.45 *μ*m filter. Calvarial osteoblasts harvested from *Rosa26^Cas9/Cas9^* pups were infected for 24 hours and recovered for 24h in regular media before further experiments.

### Mass spectrometry

Calvarial osteoblasts were cultured in 6cm plates until confluency before sample preparation for mass spectrometry. For glucose, glutamine and proline tracing experiments, naïve or differentiated calvarial cells were cultured in aMEM (Genaxxon) containing 0.3mM [U-^13^C]-Proline (Cambridge), 2mM [U-^13^C]-Glutamine (Cambridge) or 5.6mM [1,2-^13^C]-Glucose (SigmaAldrich) for 24h or 72h. The labeling was terminated with ice cold PBS and cells were scrapped with -20°C 80% methanol on dry ice. 20nmol norvaline was added into each methanol extract as internal control, followed by centrifuge at 10000 x g for 15 minutes. Supernatants were processed and analyzed by the Metabolomics Facility at the Children’s Medical Center Research Institute at UT Southwestern. For tracing experiments into protein, cells were labeled for 0, 12, 24 or 72 hours. Cell were then scrapped in 1M perchloric acid. The protein pellet was washed with 70% ethanol three times. The pellet was then incubated with 1mL of 6M HCl at 110 °C for 18 hours to hydrolyze the proteins. 1mL of chloroform was then added to each sample followed by centrifuge at 400x g for 10 minutes. Supernatants were taken for further preparation. The supernatant was dried by N2 gas at 37°C. GC-MS method for small polar metabolites assay used in this study was adapted from Wang et al. (2018). The dried residues were resuspended in 25**μ**L methoxylamine hydrochloride (2%(w/v) in pyridine) and incubated at 40°C for 90 minutes. 35 *μ*L of MTBSTFA + 1% TBDMS was then added, followed by 30-minute incubation at 60°C. The supernatants from proline tracing experiments were dried by N2 gas at 37°C followed by resuspension in 50 *μ*L of MTBSTFA + 1% TBDMS incubated at 60 °C for 30 minutes. The derivatized sampled were centrifuged for 5 minutes at 10000x g force. Supernatant from each sample was transferred to GC vials for analysis. 1*μ*L of each sample was injected in split or splitless mode depending on analyte of interest. GC oven temperature was set at 80 °C for 2 minutes, increased to 280 °C at a rate of 7 °C/min, and then kept at 280 °C for a total run time of 40 minutes.

GC-MS analysis was performed on an Agilent 7890B GC system equipped with a HP-5MS capillary column (30 m, 0.25 mm i.d., 0.25 mm-phase thickness; Agilent J&W Scientific), connected to an Agilent 5977A Mass Spectrometer operating under ionization by electron impact (Meister, 1975) at 70 eV. Helium flow was maintained at 1 mL/min. The source temperature was maintained at 230 °C, the MS quad temperature at 150 °C, the interface temperature at 280 °C, and the inlet temperature at 250 °C. Mass spectra were recorded in selected ion monitoring (SIM) mode with 4 ms dwell time.

### Amino acid uptake assay

Amino acid uptake assays were performed as previously described (Shen & Karner, 2021). Cells were first washed three times with PBS and incubated with Krebs Ringer Hepes (KRH) (120mM NaCl, 5mM KCl, 2mM CaCl_2_, 1mM MgCl_2_, 25mM NaHCO_3_, 5mM HEPES, 1mM D-Glucose) with 4µCi/mL L-[2,3,4-^3^H]-Proline (PerkinElmer NET323250UC), L-[3,4-^3^H]-Glutamine (PerkinElmer NET551250UC), L-[2,3-^3^H]-Alanine (PerkinElmer NET348250UC), L-[1,2-^14^C]-Alanine (PerkinElmer NEC266E050UC), L-[^3^H(G)]-Serine (PerkinElmer NET248250UC), L-[^14^C(U)]-Glycine (PerkinElmer NEC276E050UC), or L-[3,4-^3^H]-Glutamate (PerkinElmer NET490001MC) for 5 minutes at 37°C. Uptake and metabolism were terminated with ice cold KRH and the cells were scraped with 1% SDS. Cell lysates were combined with 8mL Ultima Gold scintillation cocktail (PerkinElmer 6013329) and CPM was measured using Beckman LS6500 Scintillation counter. Newborn mouse humeri and femurs were used for *ex vivo* amino acid uptake acid. Extemporaneous and cartilaginous tissues were removed from the bones and counter lateral parts were harvested and boiled for normalization. Bones were then incubated with KRH containing radiolabeled amino acids for 30min at 37°C. Uptake and metabolism were terminated by ice cold KRH. Samples were homogenized in RIPA lysis buffer (50 mM Tris (pH 7.4), 15 mM NaCl, 0.5% NP-40, 0.1% SDS, 0.1% sodium deoxycholate) followed by sonication using an Ultrasonic Processor (VCX130) (Amplitude: 35%, Pulse 1s, Duration: 10s) and centrifugation. Supernatant from each sample was combined with 8mL scintillation cocktail and CPM was measured using Beckman LS6500 Scintillation counter. Radioactivity was normalized with the boiled contralateral bones.

### Proline incorporation assay

Cells were incubated with KRH supplemented with 4µCi/mL L-[2,3,4-^3^H]-Proline for three hours. Cells were lysed with RIPA and followed by centrifugation. Protein is precipitated with TCA and resuspended using 1mL 1M NaOH. 200uL of the dissolved sample was saved for radioactivity reading later as the total proteins. The rest of each sample was split into two: one was treated with 15mg Collagenase P and 60mM HEPES to digest collagens and the other with only 60mM HEPES as the baseline control. Samples were incubated at 37°C for 3hours. After incubation, residual proteins and Collagenase P was precipitated using TCA followed by centrifugation. Supernatant from each sample was combined with 8mL scintillation cocktail and CPM was measured using Beckman LS6500 Scintillation counter. Radioactivity for collagen incorporation was normalized with 60mM HEPES treated the baseline control.

### RNA isolation and qPCR

Total RNA was harvested from calvarial osteoblasts using TRIZOL and purified by mixing with chloroform. 500ng of total RNA was used for reverse transcription by IScript cDNA synthesis kit (Bio-Rad). SsoAdvanced Universal SYBR Green Supermix (Bio-Rad) was used for qPCR with primers used at 0.1*μ*M (listed in Table S3). Technical and biological triplicates were performed using a 96-well plate on an ABI QuantStudio 3. The PCR program was set as 95℃ for 3min followed by 40 cycles of 95℃ for 10s and 60℃ for 30s. *Actb* mRNA level was used to normalize expression of genes of interest and relative expression was calculated using the 2^-(ΔΔCt)^ method. PCR efficiency was optimized and melting curve analyses of products were performed to ensure reaction specificity.

### RNAseq

RNA sequencing was performed in biological triplicate by the Duke University Center for Genomic and Computational Biology Sequencing and Genomic Technology Shared Resource on 10mg of RNA isolated from primary calvarial cells cultured in either growth or osteogenic media for 7 days. RNA-seq data was processed using the TrimGalore toolkit1 which employs Cutadapt2 to trim low-quality bases and Illumina sequencing adapters from the 3’ end of the reads. Only reads that were 20nt or longer after trimming were kept for further analysis. Reads were mapped to the GRCm38v68 version of the mouse genome and transcriptome3 using the STAR RNA-seq alignment tool4. Reads were kept for subsequent analysis if they mapped to a single genomic location. Gene counts were compiled using the HTSeq tool5. Only genes that had at least 10 reads in any given library were used in subsequent analysis. Normalization and differential expression were carried out using the DESeq26 Bioconductor7 package with the R statistical programming environment8. The false discovery rate was calculated to control for multiple hypothesis testing. Gene set enrichment analysis9 was performed to identify differentially regulated pathways and gene ontology terms for each of the comparisons performed.

### Western blotting

Calvarial osteoblasts were scraped in RIPA lysis buffer with cOmplete protease inhibitor and PhosSTOP cocktail tablets (Roche). Protein concentration was determined by BCA protein assay kit (Thermo). Protein (6-20 **μ** g) was loaded on 4%-15% or 12% polyacrylamide gel and transferred onto Immuno-Blot PVDF membrane. The membranes were blocked for 1 hour at room temperature in 5% milk powder in TBS with 0.1% Tween (TBST) and then incubated at 4°C with the primary antibody overnight. Primary antibodies were used at 1:1000 to detect proteins, listed as follows: Anti-SNAT2 (AB_2050321), Anti-P-S240/244 S6 (AB_331682), Anti-S6 (AB_331355), Anti-P-S51 Eif2a (AB_2096481), Anti-Eif2a (AB_10692650), Anti-Col1a1 (AB_1672342), Anti-Runx2 (AB_10949892), Anti-β−actin (AB_330288), Anti-Smad1 (AB_2107780), Anti-4E-BP1 (AB_2097841), Anti-ATF4 (AB_2058752), Anti-mTOR (AB_2105622), Anti-Akt (AB_329827), Anti-Erk (AB_390779), Anti-eEF2 (AB_10693546), Anti-Phgdh (AB_2750870), Anti-α-Tubulin (AB_2619646), Anti-Osx (AB_2895257). Membranes were then incubated at room temperature with Anti-Rabbit IgG (AB_2099233) or Anti-Mouse IgG, HRP-linked Antibody (AB_330924) at 1:2000 for 1 hour at room temperature. Immunoblots were next developed by enhanced chemiluminescence (Clarity Substrate Kit or SuperSignal West Femto substrate). Each experiment was repeated with at least three independently prepared protein extractions. Densitometry was performed for quantification for each blot.

### Amino acid proportion and amino acid demand prediction analysis

Amino acid sequences of proteins (Mus_musculus.GRCm38.pep.all.fa) were retrieved from Ensembl (https://uswest.ensembl.org/info/data/ftp/index.html). Amino acid proportion was calculated based on the amino acid sequences of specific proteins (RUNX2, COL1A1, OSX and OCN) and proteins associated with different GO terms. mRNA expression of genes in undifferentiated and differentiated osteoblasts were obtained from transcriptomic analysis. Top 500 induced and suppressed genes from differentiated osteoblasts were selected for the calculation of proline proportion. For amino acid demand prediction, amino acid proportion and mRNA expression were merged using *Gene.stable.ID* as the bridge. 75 unmatched proteins were excluded from a total of 49665 proteins. To predict the amino acid demand change, changes in mRNA expression was assumed to be proportional to changes in protein translation. Based on this, the change of amino acid demand in each protein is proportional to mRNA expression change:

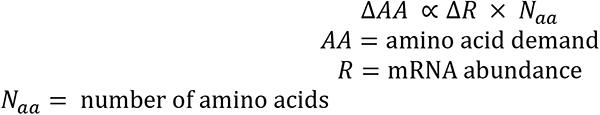

To summarize the overall change of amino acid demand during osteoblast differentiation:

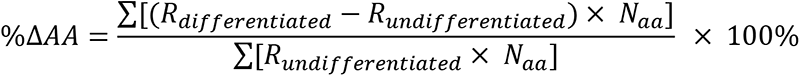

### tRNA aminoacylation assay

The method is adapted from (Loayza-Puch et al., 2016; Saikia et al., 2016). Purified RNA was resuspended in 30mM NaOAc/HOAc (pH 4.5). RNA was divided into two parts (2**μ**g each): one was oxidized with 50mM NaIO_4_ in 100mM NaOAc/HOAc (pH 4.5) and the other was treated with 50mM NaCl in NaOAc/HOAc (pH 4.5) for 15 min at room temperature. Samples were quenched with 100mM glucose for 5min at room temperature, followed by desalting using G50 columns and precipitation using ethanol. tRNA was then deacylated in 50mM Tris-HCl (pH 9) for 30min at 37℃, followed by another ethanol precipitation. RNA (400ng) was then ligated the 3′adaptor (5′-/5rApp/TGGAATTCTCGGGTGCCAAGG/3ddC/-3′) using T4 RNA ligase 2 (NEB) for 4 h at 37°C. 1**μ**g RNA was then reverse transcribed using SuperScript III first strand synthesis system with the primer (GCCTTGGCACCCGAGAATTCCA) following the manufacturer’s instruction. Relative charging level was calculated by qRT-PCR using tRNA-specific primers stated in Table S2.

### Flow Cytometry

Flow cytometry was used to analyze EdU incorporation and cell viability in calvarial osteoblasts. EdU incorporation was performed using Click-iT™ EdU Alexa Fluor™ 488 Flow Cytometry Assay Kit. Cell were incubated with EdU (5-ethynyl-2′-deoxyuridine, 10 **μ**M) for 24 hours. Cells were then trypsinized, fixed, permeabilized and incubated with Click-iT reaction cocktail for 30 minutes according to manufacturer’s instructions. Cell viability was analyzed using the Cell Meter™ APC-Annexin V Binding Apoptosis Assay Kit (Cat# 22837). Cells were trypsinized and incubated with APC-Annexin V conjugate and propidium iodide for 30min. Cells were all resuspended in 500**μ**L PBS and analyzed using FACSCanto II flow cytometer (BD Biosciences). Data were analyzed and evaluated using FlowJo (v.11).

### Quantification and Statistical analysis

Statistical analyses were performed using either Graphpad Prism 6 or R software. One-way ANOVA or unpaired 2-tailed Student’s t-test were used to determine statistical significance as indicated in the text. All data are shown as mean values ± SD or SEM as indicated. p<0.05 is considered as statistically significant. Sample size (n) and other statistical parameters are included in the figure legends. Experiments were repeated on a minimum of 3 independent samples unless otherwise noted.

## Acknowledgements

The authors thank Drs. Thomas Carroll and Guoli Hu for critical comments on this manuscript. This work was supported by the National Institute of Arthritis and Musculoskeletal and Skin Diseases (NIAMS) grants (AR076325 and AR071967) to C.M.K.

## Author Contributions

Conceptualization, C.M.K; Investigation, L.S., Y.Y., Y.Z. S.PM., G.Z., and C.M.K.; Writing – Original Draft, L.S.; Writing – Review & Editing, G.Z. and C.M.K.; Supervision, C.M.K.

## Supplemental Figure Legends

**Figure 1 Supplement 1.**
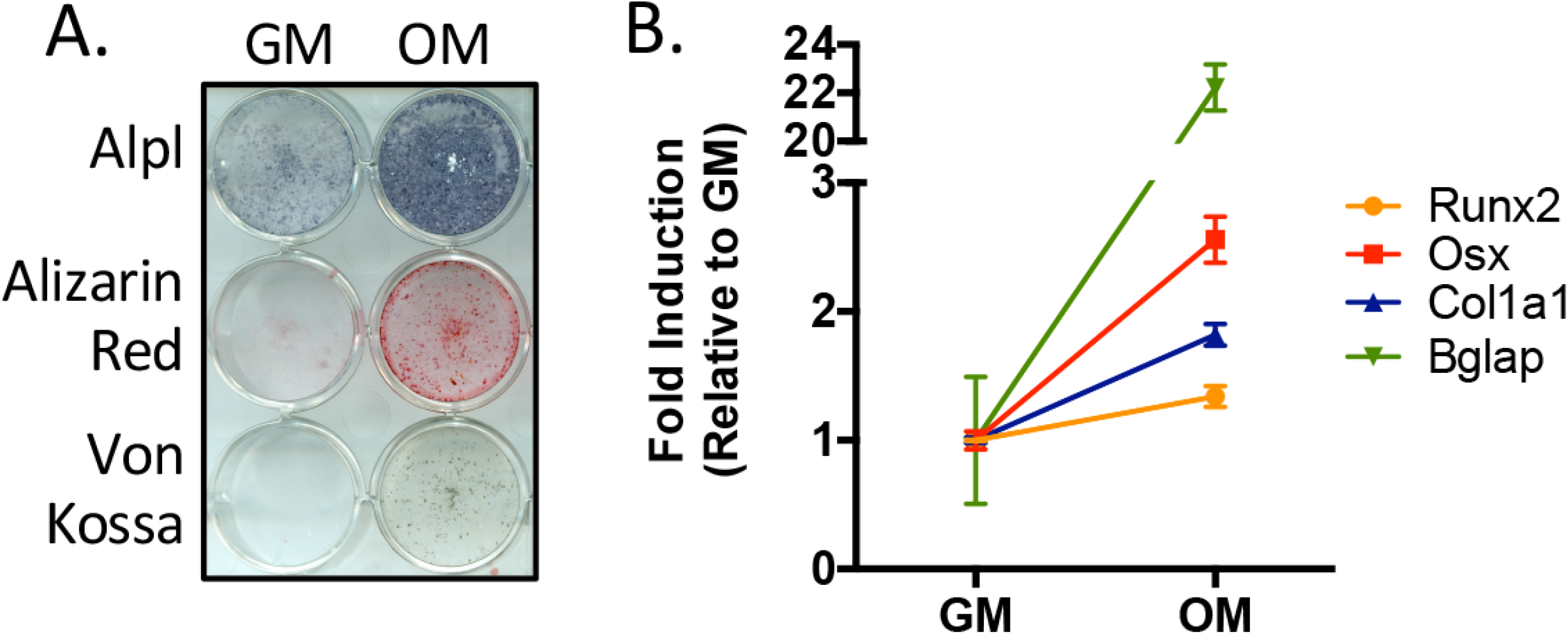
**(A)** Functional assays of calvarial cells cultured in growth media (GM) or osteogenic medium (OM) for 10 days. **(B)** qRT-PCR analysis of osteogenic marker genes *Runx2*, *Sp7* (OSX), *Col1a1* and *Bglap* in calvarial cells cultured in GM or OM for 7 days (n=3). Error bars depict SD.

**Figure 2 Supplement 1.**
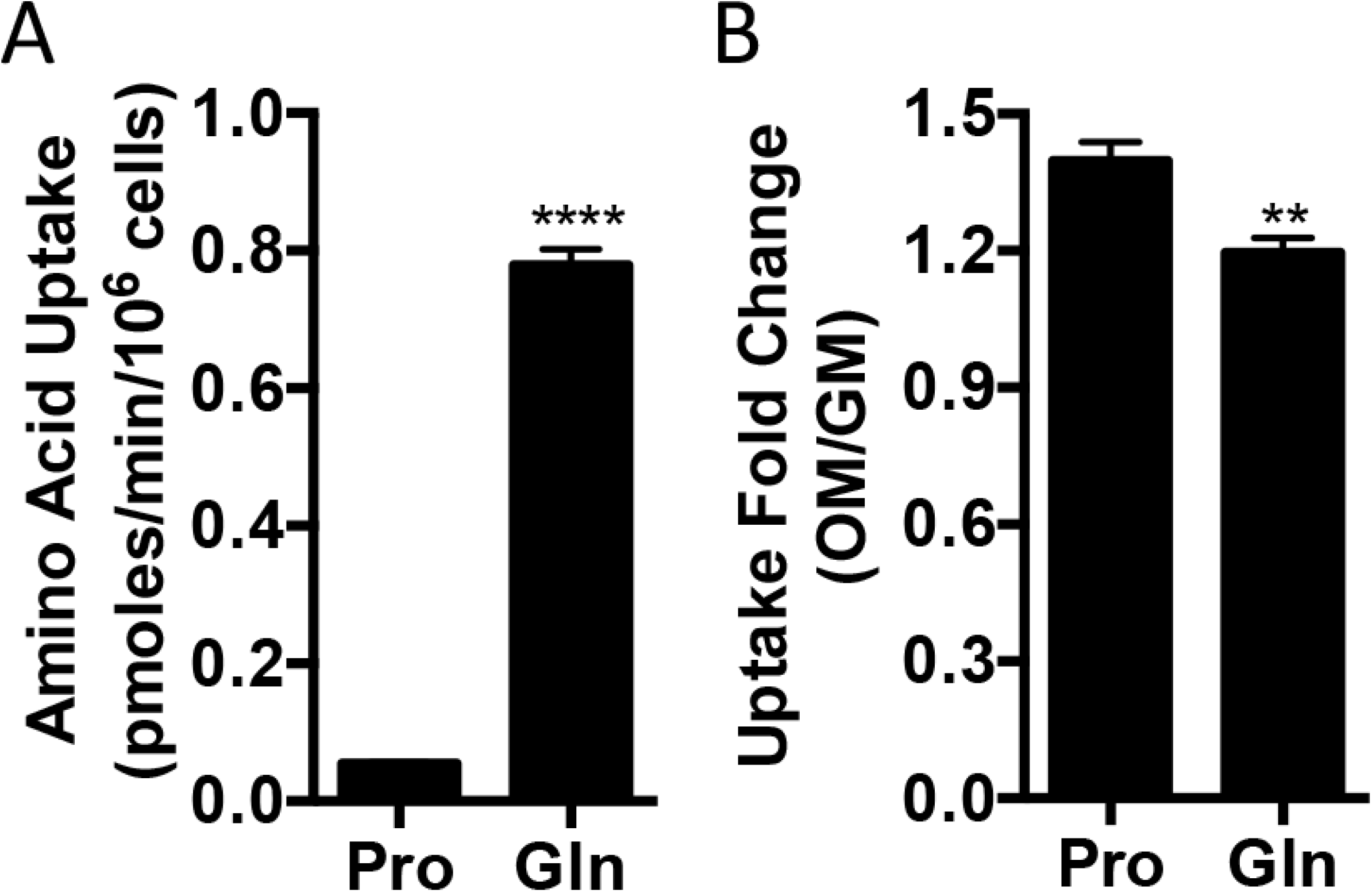
**(A,B)** Radiolabeled proline or glutamine uptake assay performed in naïve calvarial osteoblasts **(A)** or after 7 days of osteoblast differentiation **(B)**. Error bars depict SD. ** p≤ 0.005, **** p≤ 0.00005. by unpaired 2-tailed Student’s t-test.

**Figure 3 Supplement 1.**
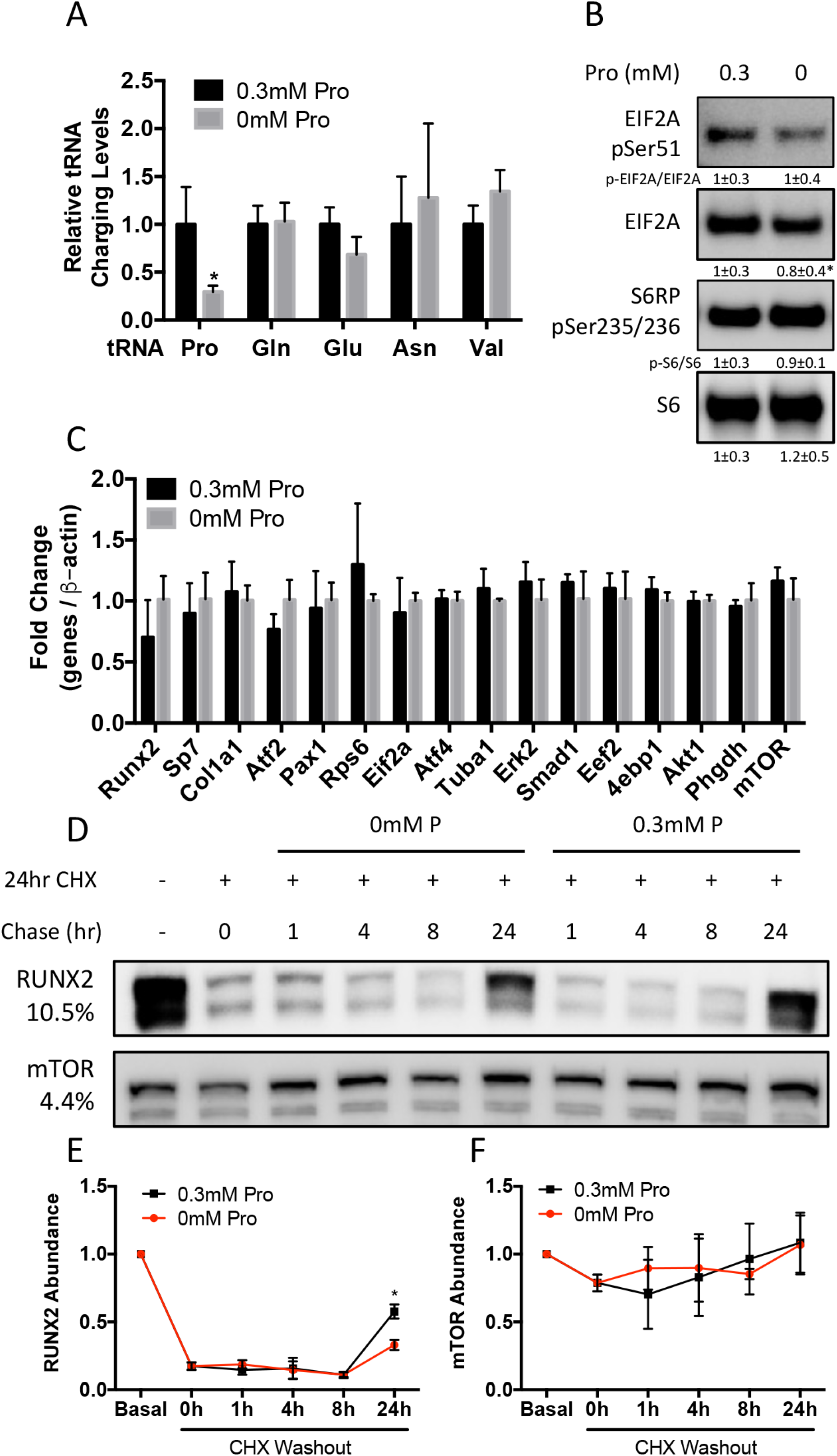
**(A)** Effect of 48-hour proline withdrawal on tRNA aminoacylation (n=3). **(B)** Western blot analysis of naïve calvarial cells cultured in 0.3mM or 0mM Pro for 48 hours (n=3). Protein expression normalized to total protein. Phosphorylation normalized to total protein. Fold change ± SD for three independent experiments. **(C)** qRT-PCR analysis of the effect of 48-hour proline withdrawal on gene expression in calvarial cells (n=3). **(D-F)** The effect of proline availability on the synthesis of select proteins. CHX – cycloheximide. Fold change ± SD for three independent experiments. Error bars depict SD. * p≤ 0.05. by unpaired 2-tailed Student’s t-test.

**Figure 4 Supplement 1.**
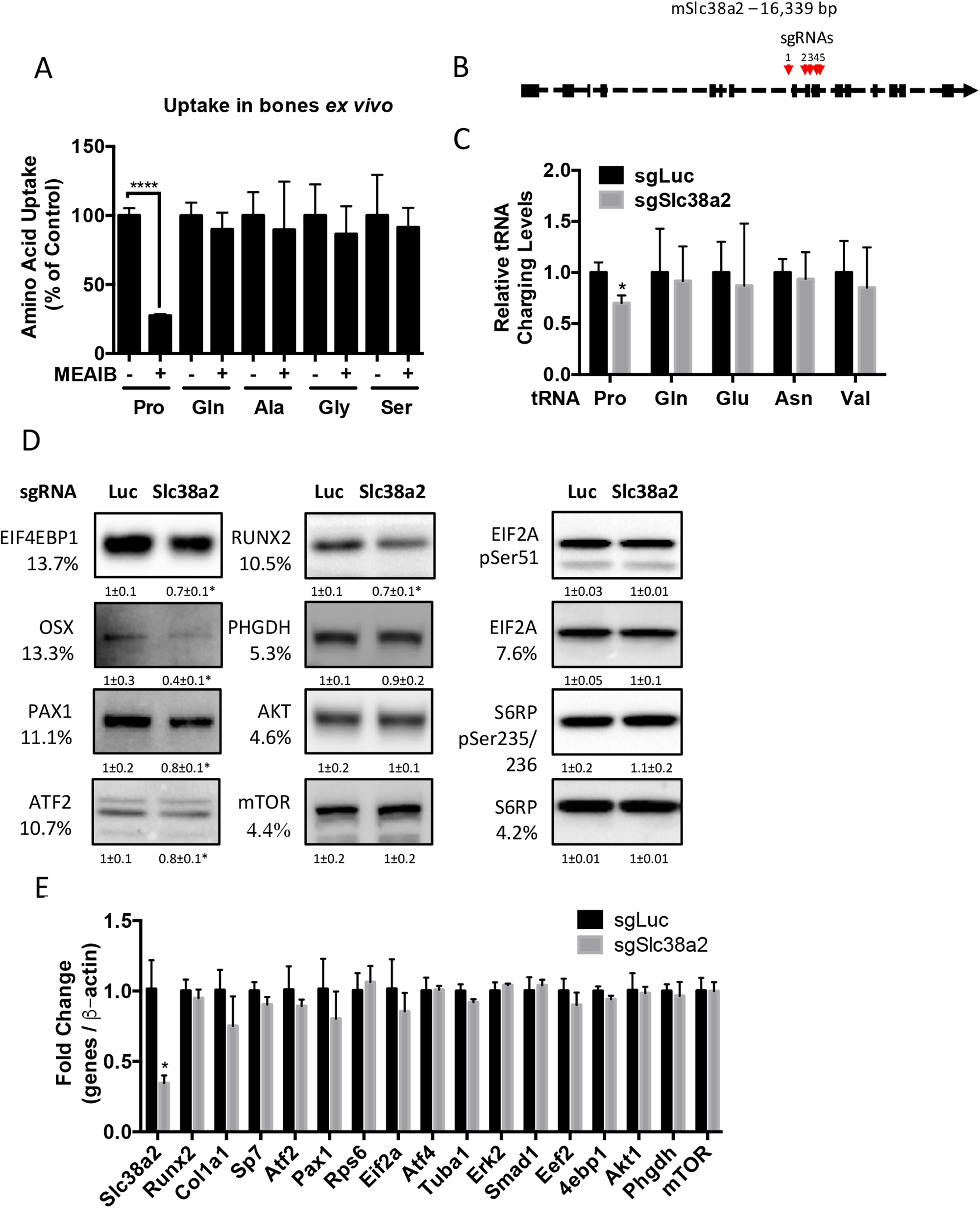
**(A)** Graphical depiction of the effects of 5mM MEAIB on amino acid uptake in humeri or femurs isolated from P3 mice (n=3). **(B)** Schematic depicting *Slc38a2* Crispr targeting strategy. **(C-E)** Effect of *Slc38a2* targeting on tRNA aminoacylation (n=3) **(C),** protein expression (n=3) **(D)** or mRNA expression (n=3) **(E)**. In all blots, the percent proline composition is noted under the protein name. Protein expression normalized to total protein. Phosphorylation normalized to total protein. Fold change ± SD for three independent experiments. * p≤ 0.05, **** p≤ 0.00005. by unpaired 2-tailed Student’s t-test.

**Figure 5 Supplement 1.**
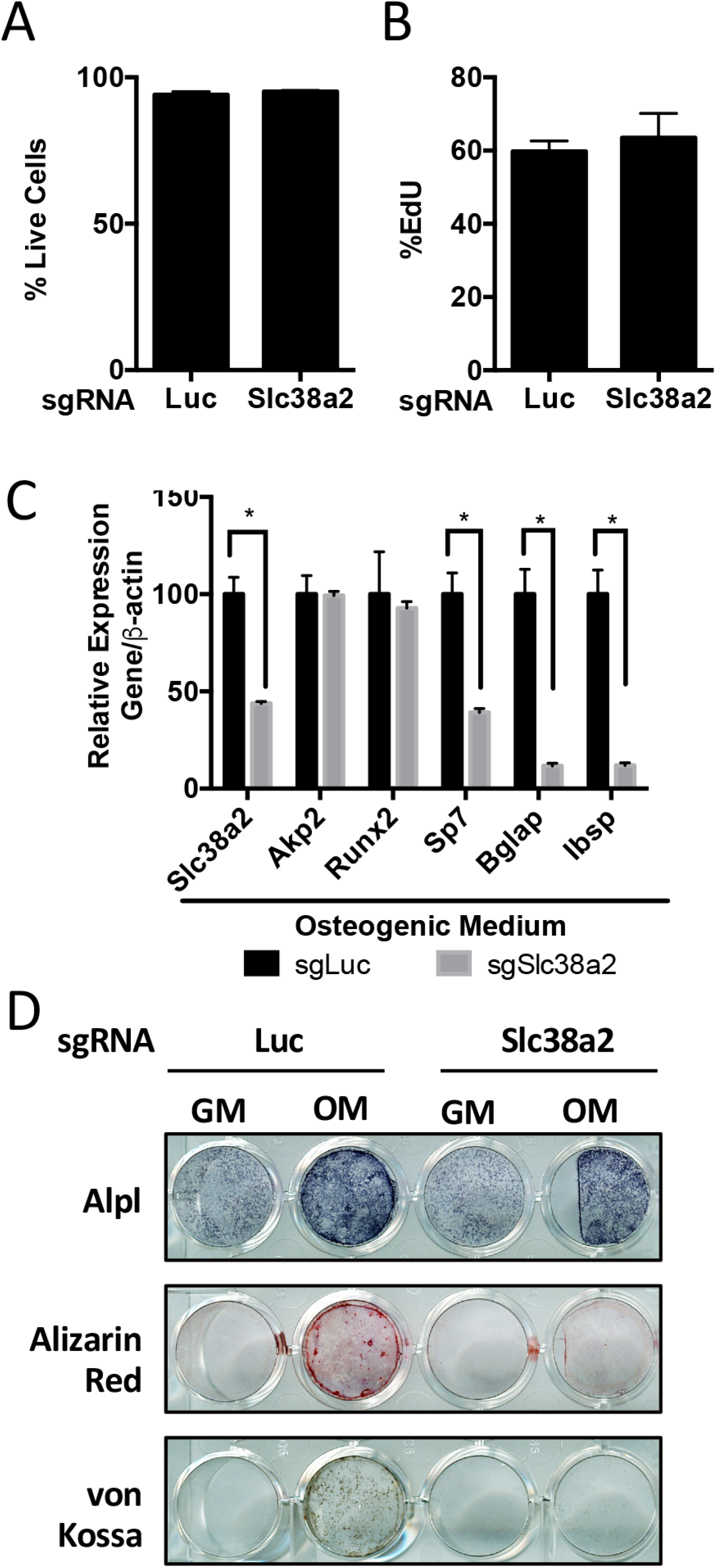
**(A-D)** Effect of *Slc38a2* targeting on cell viability (n=3) **(A)**, EdU incorporation (n=3) **(B)**, mRNA expression (n=3) by qPCR analysis **(C)**, or functional assays **(D)** in calvarial cells cultured in growth media (GM) or osteogenic medium (OM) for 7 or 10 days. Error bars depict SD. * p≤ 0.05. by unpaired 2-tailed Student’s t-test.

**Figure 5 Supplement 2.**
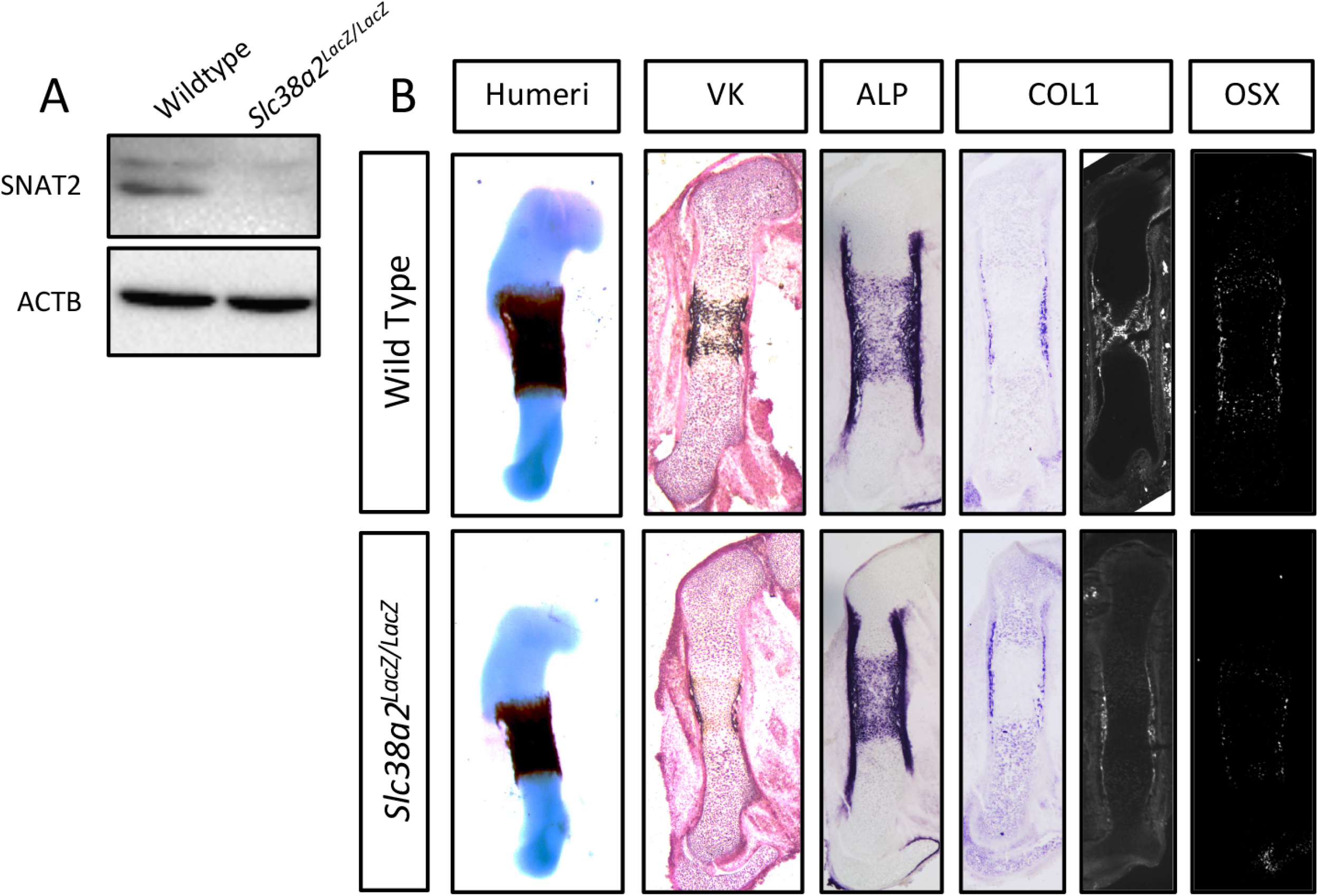
**(A)** Western blot analysis of SNAT2 expression in femurs from *Slc38a2^LacZ/LacZ^* or wildtype controls. SNAT2 proteins normalized to ACTB. **(B)** Representative images of humerus skeletal preparations, von Kossa staining, alkaline phosphatase (ALPL) staining, *in situ* hybridization for *Col1a1*, or immunofluorescence staining for OSX and COL1A1 on femur sections of E15.5 *Slc38a2^LacZ/LacZ^* or wildtype controls littermate controls (n=3 animals).

**Figure 5 Supplement 3.**
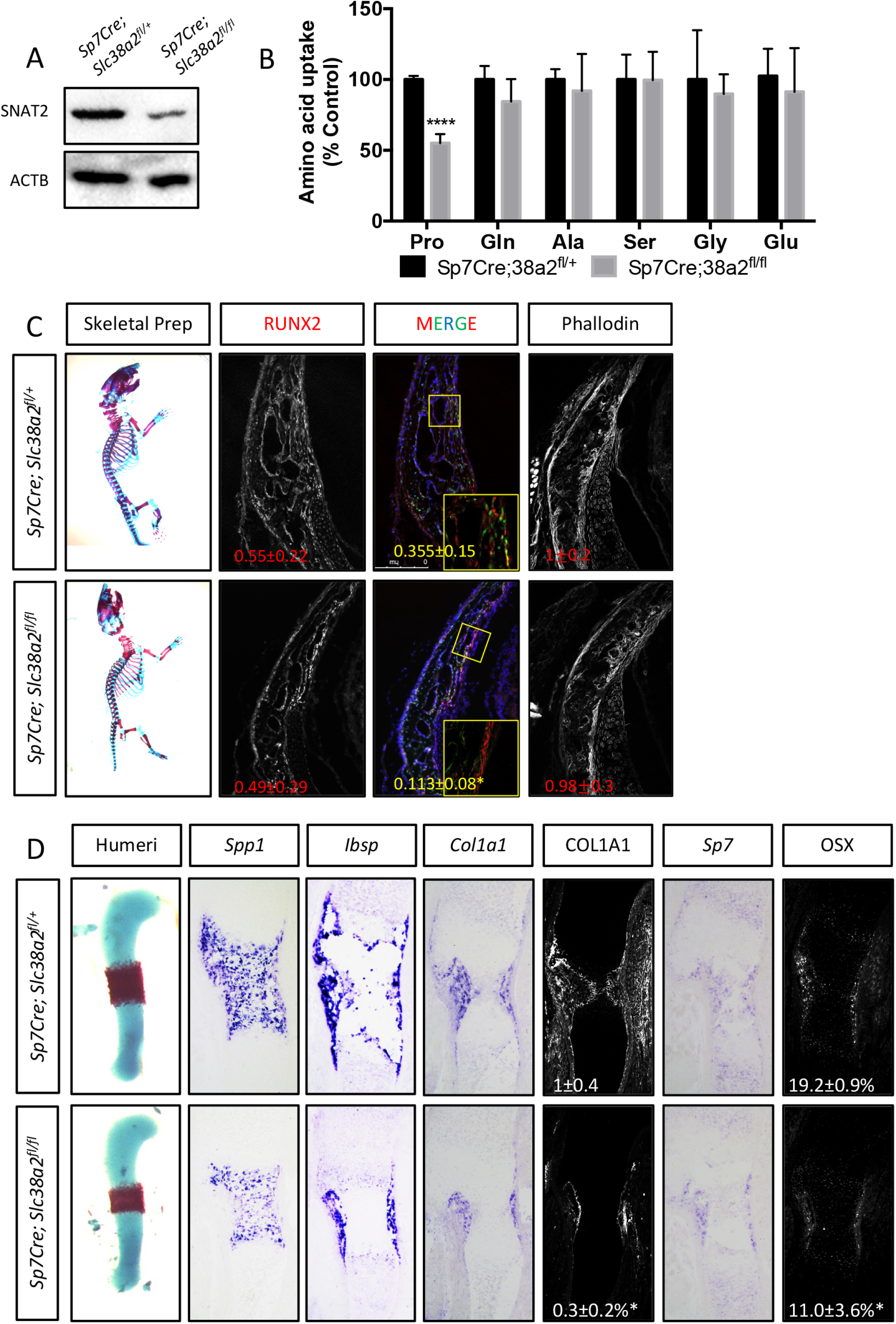
**(A)** Western blot analysis of SNAT2 expression in femurs from *Sp7Cre;Slc38a2^fl/fl^* or *Sp7Cre;Slc38a2^fl/+^* littermate controls. SNAT2 normalized to ACTB. **(B)** Evaluation of amino acid uptake in femurs isolated from newborn *Sp7Cre;Slc38a2^fl/fl^* or *Sp7Cre;Slc38a2^fl/+^* littermate controls (n=5). **(C)** Skeletal preparations of newborn *Slc38a2^LacZ/LacZ^* or wildtype controls (n=5), or *Sp7Cre;Slc38a2^fl/fl^* or *Sp7Cre;Slc38a2^fl/+^* littermate controls (n=5). Phalloidin staining or immunofluorescence staining for RUNX2 on P0 *Sp7Cre;Slc38a2^fl/fl^* or *Sp7Cre;Slc38a2^fl/+^* calvariae (n=5). **(D)** Representative images of *in situ* hybridization for *Spp1, Ibsp, Col1a1*, and *Sp7*, or immunofluorescence staining for OSX and COL1A1 on humerus sections from E15.5 *Sp7Cre;Slc38a2^fl/fl^* or *Sp7Cre;Slc38a2^fl/+^* littermate controls (n=5 animals). Fold change ± SD. Error bar depicts SD. *p≤0.05, ****p≤0.00005. by paired 2-tailed Student’s t-test.

**Table S1.**
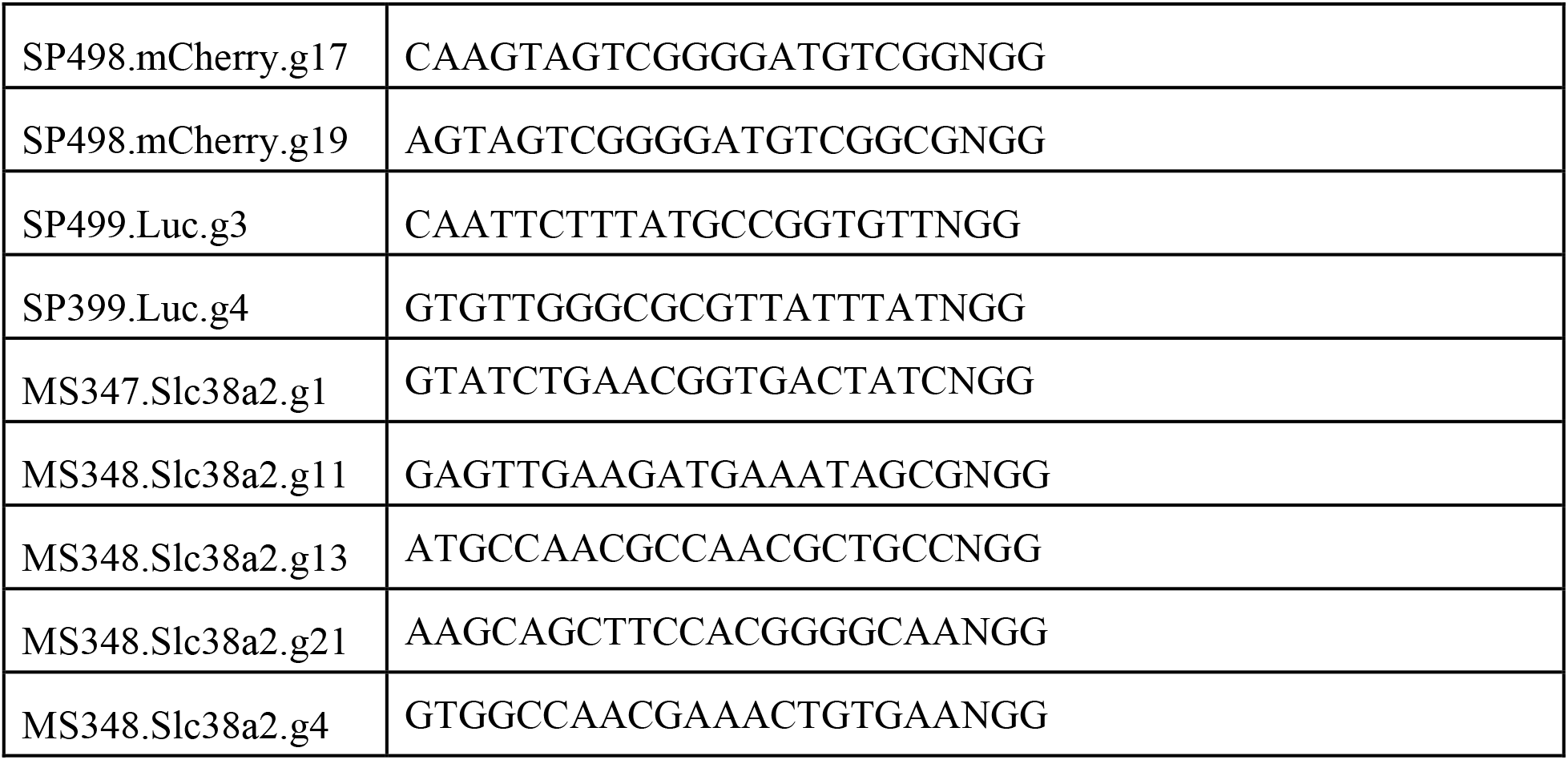
sgRNA protospacer sequence.

**Table S2.**
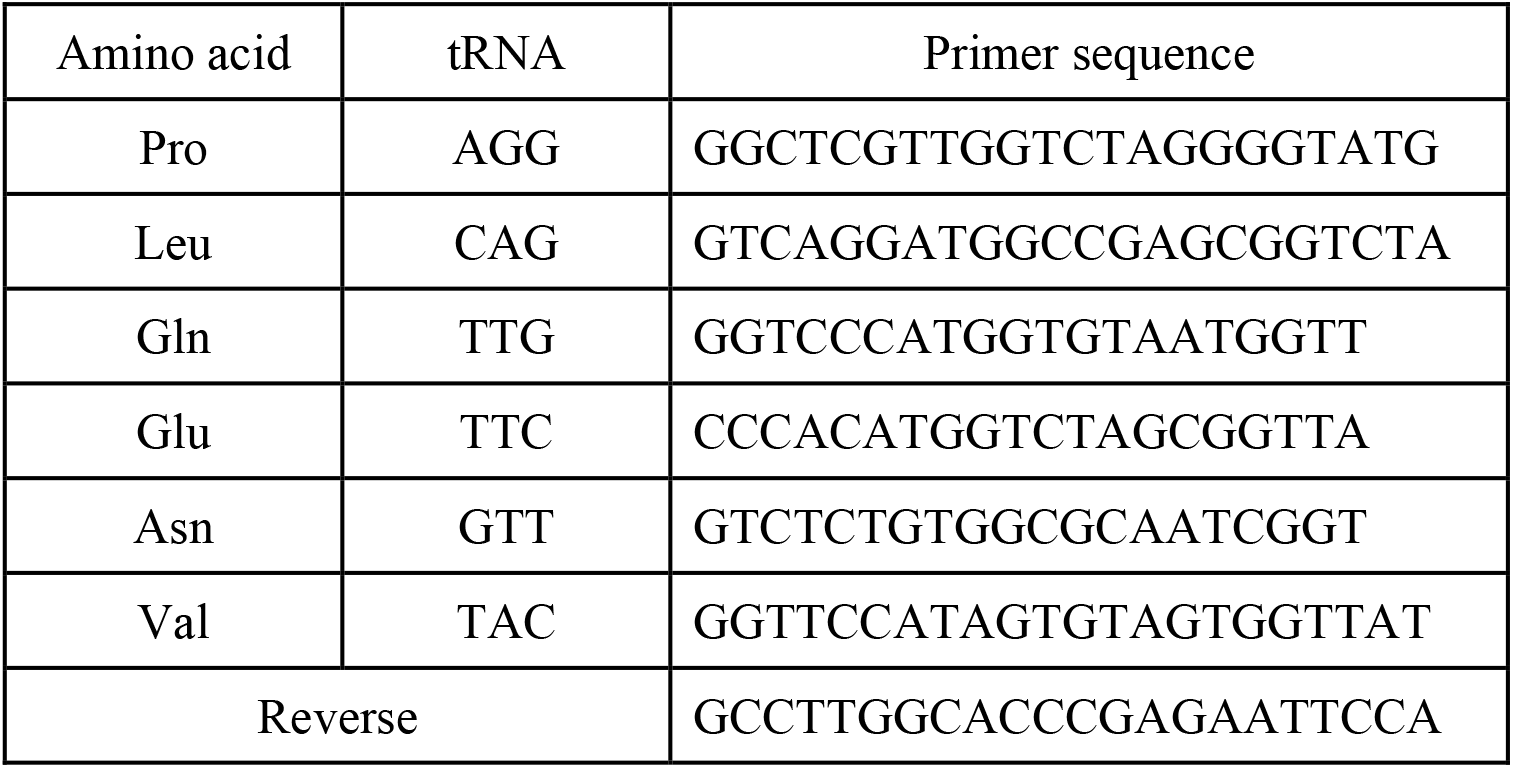
RT-PCR primer sequences for tRNA charging.

**Table S3.**
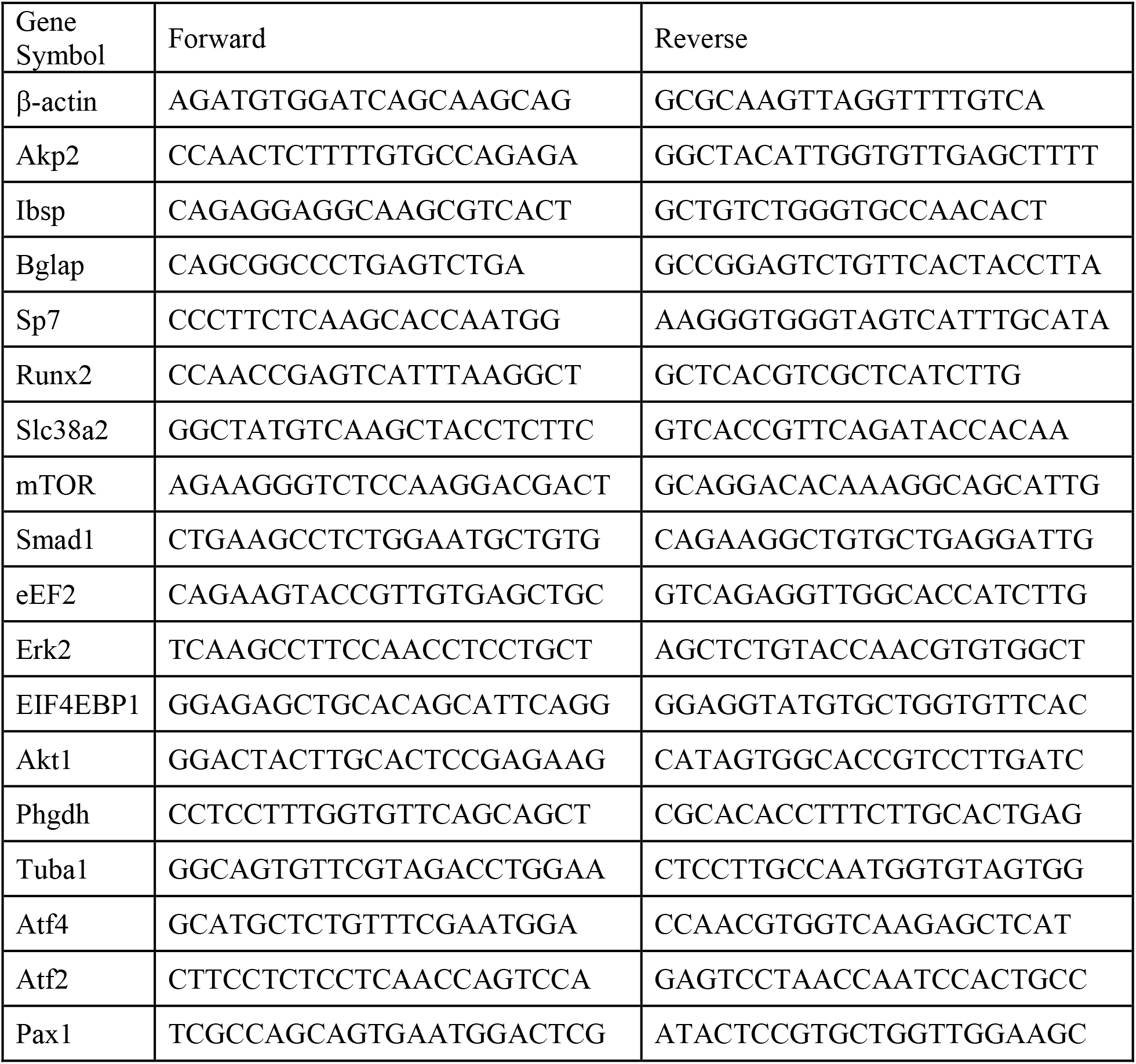
RT-PCR primer sequences.

## References

Adamson, L. F., & Ingbar, S. H. (1967). Some properties of the stimulatory effect of thyroid hormones on amino acid transport by embryonic chick bone. Endocrinology, 81(6), 1372–1378. doi:10.1210/endo-81-6-1372

Alves, R. D. A. M., Eijken, M., Swagemakers, S., Chiba, H., Titulaer, M. K., Burgers, P. C., … van Leeuwen, J. P. T.M. (2010). Proteomic Analysis of Human Osteoblastic Cells: Relevant Proteins and Functional Categories for Differentiation. Journal of Proteome Research, 9(9), 4688–4700. doi:10.1021/pr100400d

Baek, W. Y., Park, S. Y., Kim, Y. H., Lee, M. A., Kwon, T. H., Park, K. M., … Kim, J.E. (2013). Persistent low level of osterix accelerates interleukin-6 production and impairs regeneration after tissue injury. PLoS One, 8(7), e69859. doi:10.1371/journal.pone.0069859

Bardai, G., Lemyre, E., Moffatt, P., Palomo, T., Glorieux, F. H., Tung, J., … Rauch, F. (2016). Osteogenesis Imperfecta Type I Caused by COL1A1 Deletions. Calcified Tissue International, 98(1), 76–84. doi:10.1007/s00223-015-0066-6

Baum, B. J., & Shteyer, A. (1987). Characteristics of a neutral amino acid transport system (system A) in osteoblastic rat osteosarcoma cells. Exp Cell Res, 169(2), 453–457. doi:10.1016/0014-4827(87)90205-9

Ben Amor, I. M., Roughley, P., Glorieux, F. H., & Rauch, F. (2013). Skeletal clinical characteristics of osteogenesis imperfecta caused by haploinsufficiency mutations in COL1A1. Journal of Bone and Mineral Research, 28(9), 2001–2007. doi:https://doi.org/10.1002/jbmr.1942

Berendsen, A. D., & Olsen, B. R. (2015). Bone development. Bone, 80, 14–18. doi:10.1016/j.bone.2015.04.035

Bianco, P., Fisher, L. W., Young, M. F., Termine, J. D., & Robey, P. G. (1991). Expression of bone sialoprotein (BSP) in developing human tissues. Calcif Tissue Int, 49(6), 421–426. doi:10.1007/BF02555854

Bröer, A., Rahimi, F., & Bröer, S. (2016). Deletion of amino acid transporter ASCT2 (SLC1A5) reveals an essential role for transporters SNAT1 (SLC38A1) and SNAT2 (SLC38A2) to sustain glutaminolysis in cancer cells. Journal of biological chemistry, 291(25), 13194–13205.

Buttgereit, F., & Brand, M. D. (1995). A hierarchy of ATP-consuming processes in mammalian cells. Biochem J, 312 (Pt 1)(Pt 1), 163–167. doi:10.1042/bj3120163

Chen, J., & Long, F. (2018). mTOR signaling in skeletal development and disease. Bone Research, 6(1), 1. doi:10.1038/s41413-017-0004-5

Choi, J. Y., Pratap, J., Javed, A., Zaidi, S. K., Xing, L., Balint, E., … Stein, G.S. (2001). Subnuclear targeting of Runx/Cbfa/AML factors is essential for tissue-specific differentiation during embryonic development. Proc Natl Acad Sci U S A, 98(15), 8650–8655. doi:10.1073/pnas.151236498

Ducy, P., Desbois, C., Boyce, B., Pinero, G., Story, B., Dunstan, C., … Karsenty, G. (1996). Increased bone formation in osteocalcin-deficient mice. Nature, 382(6590), 448–452. doi:10.1038/382448a0

Ducy, P., Zhang, R., Geoffroy, V., Ridall, A. L., & Karsenty, G. (1997). Osf2/Cbfa1: a transcriptional activator of osteoblast differentiation. Cell, 89(5), 747–754. doi:10.1016/s0092-8674(00)80257-3

Elefteriou, F., Benson, M. D., Sowa, H., Starbuck, M., Liu, X., Ron, D., … Karsenty, G. (2006). ATF4 mediation of NF1 functions in osteoblast reveals a nutritional basis for congenital skeletal dysplasiae. Cell Metab, 4(6), 441–451. doi:10.1016/j.cmet.2006.10.010

Esen, E., Chen, J., Karner, C. M., Okunade, A. L., Patterson, B. W., & Long, F. (2013). WNT-LRP5 signaling induces Warburg effect through mTORC2 activation during osteoblast differentiation. Cell Metab, 17(5), 745–755. doi:10.1016/j.cmet.2013.03.017

Finerman, G. A., & Rosenberg, L. E. (1966). Amino acid transport in bone. Evidence for separate transport systems for neutral amino and imino acids. J Biol Chem, 241(7), 1487–1493. Retrieved from https://www.ncbi.nlm.nih.gov/pubmed/5946610

Grant, M. E., & Prockop, D. J. (1972). The Biosynthesis of Collagen. New England Journal of Medicine, 286(4), 194–199. doi:10.1056/NEJM197201272860406

Grewal, S., Defamie, N., Zhang, X., De Gois, S., Shawki, A., Mackenzie, B., … Erickson, J.D. (2009). SNAT2 amino acid transporter is regulated by amino acids of the SLC6 gamma-aminobutyric acid transporter subfamily in neocortical neurons and may play no role in delivering glutamine for glutamatergic transmission. The Journal of biological chemistry, 284(17), 11224–11236. doi:10.1074/jbc.M806470200

Guntur, A. R., & Rosen, C. J. (2012). Bone as an endocrine organ. Endocrine practice: official journal of the American College of Endocrinology and the American Association of Clinical Endocrinologists, 18(5), 758–762. doi:10.4158/EP12141.RA

Hahn, T. J., Downing, S. J., & Phang, J. M. (1969). Amino acid transport in adult diaphyseal bone: contrast with amino acid transport mechanisms in fetal membranous bone. Biochim Biophys Acta, 183(1), 194–203. doi:10.1016/0005-2736(69)90143-6

Hahn, T. J., Downing, S. J., & Phang, J. M. (1971). Insulin effect on amino acid transport in bone: dependence on protein synthesis and Na+. American Journal of Physiology-Legacy Content, 220(6), 1717–1723. doi:10.1152/ajplegacy.1971.220.6.1717

Hoffmann, T. M., Cwiklinski, E., Shah, D. S., Stretton, C., Hyde, R., Taylor, P. M., & Hundal, H. S. (2018). Effects of Sodium and Amino Acid Substrate Availability upon the Expression and Stability of the SNAT2 (SLC38A2) Amino Acid Transporter. Frontiers in Pharmacology, 9(63). doi:10.3389/fphar.2018.00063

Hollinshead, K. E. R., Munford, H., Eales, K. L., Bardella, C., Li, C., Escribano-Gonzalez, C., … Tennant, D.A. (2018). Oncogenic IDH1 Mutations Promote Enhanced Proline Synthesis through PYCR1 to Support the Maintenance of Mitochondrial Redox Homeostasis. Cell Rep, 22(12), 3107–3114. doi:10.1016/j.celrep.2018.02.084

Hu, G., Yu, Y., Tang, Y. J., Wu, C., Long, F., & Karner, C. M. (2020). The Amino Acid Sensor Eif2ak4/GCN2 Is Required for Proliferation of Osteoblast Progenitors in Mice. Journal of Bone and Mineral Research, 35(10), 2004–2014. doi:https://doi.org/10.1002/jbmr.4091

Jagannathan-Bogdan, M., & Zon, L. I. (2013). Hematopoiesis. *Development* (Cambridge, England), 140(12), 2463–2467. doi:10.1242/dev.083147

Kandasamy, P., Gyimesi, G., Kanai, Y., & Hediger, M. A. (2018). Amino acid transporters revisited: New views in health and disease. Trends Biochem Sci, 43(10), 752–789. doi:10.1016/j.tibs.2018.05.003

Karner, C. M., Esen, E., Okunade, A. L., Patterson, B. W., & Long, F. (2015). Increased glutamine catabolism mediates bone anabolism in response to WNT signaling. J Clin Invest, 125(2), 551–562. doi:10.1172/JCI78470

Kern, B., Shen, J., Starbuck, M., & Karsenty, G. (2001). Cbfa1 contributes to the osteoblast-specific expression of type I collagen genes. J Biol Chem, 276(10), 7101–7107. doi:10.1074/jbc.M006215200

Komori, T., Yagi, H., Nomura, S., Yamaguchi, A., Sasaki, K., Deguchi, K., … Kishimoto, T. (1997). Targeted disruption of Cbfa1 results in a complete lack of bone formation owing to maturational arrest of osteoblasts. Cell, 89(5), 755–764. doi:10.1016/s0092-8674(00)80258-5

Krane, S. M. (2008). The importance of proline residues in the structure, stability and susceptibility to proteolytic degradation of collagens. Amino Acids, 35(4), 703–710. doi:10.1007/s00726-008-0073-2

Lapunzina, P., Aglan, M., Temtamy, S., Caparrós-Martín, J. A., Valencia, M., Letón, R., … Ruiz-Perez, V.L. (2010). Identification of a frameshift mutation in Osterix in a patient with recessive osteogenesis imperfecta. Am J Hum Genet, 87(1), 110–114. doi:10.1016/j.ajhg.2010.05.016

Lee, B., Thirunavukkarasu, K., Zhou, L., Pastore, L., Baldini, A., Hecht, J., … Karsenty, G. (1997). Missense mutations abolishing DNA binding of the osteoblast-specific transcription factor OSF2/CBFA1 in cleidocranial dysplasia. Nat Genet, 16(3), 307–310. doi:10.1038/ng0797-307

Lee, W.-C., Ji, X., Nissim, I., & Long, F. (2020). Malic Enzyme Couples Mitochondria with Aerobic Glycolysis in Osteoblasts. Cell Reports, 32(10), 108108. doi:https://doi.org/10.1016/j.celrep.2020.108108

Liu, W., Glunde, K., Bhujwalla, Z. M., Raman, V., Sharma, A., & Phang, J. M. (2012). Proline oxidase promotes tumor cell survival in hypoxic tumor microenvironments. Cancer Res, 72(14), 3677–3686. doi:10.1158/0008-5472.CAN-12-0080

Liu, W., Le, A., Hancock, C., Lane, A. N., Dang, C. V., Fan, T. W., & Phang, J. M. (2012). Reprogramming of proline and glutamine metabolism contributes to the proliferative and metabolic responses regulated by oncogenic transcription factor c-MYC. Proc Natl Acad Sci U S A, 109(23), 8983–8988. doi:10.1073/pnas.1203244109

Loayza-Puch, F., Rooijers, K., Buil, L. C., Zijlstra, J., Oude Vrielink, J. F., Lopes, R., … Agami, R. (2016). Tumour-specific proline vulnerability uncovered by differential ribosome codon reading. Nature, 530(7591), 490–494. doi:10.1038/nature16982

Long, F. (2012). Building strong bones: molecular regulation of the osteoblast lineage. Nature Reviews Molecular Cell Biology, 13(1), 27–38. doi:10.1038/nrm3254

Lou, Y., Javed, A., Hussain, S., Colby, J., Frederick, D., Pratap, J., … Stein, J.L. (2009). A Runx2 threshold for the cleidocranial dysplasia phenotype. Human Molecular Genetics, 18(3), 556–568. doi:10.1093/hmg/ddn383

Meyer, M. B., Benkusky, N. A., Lee, C. H., & Pike, J. W. (2014). Genomic determinants of gene regulation by 1,25-dihydroxyvitamin D3 during osteoblast-lineage cell differentiation. J Biol Chem, 289(28), 19539–19554. doi:10.1074/jbc.M114.578104

Morotti, M., Bridges, E., Valli, A., Choudhry, H., Sheldon, H., Wigfield, S., … Jones, D. (2019). Hypoxia-induced switch in SNAT2/SLC38A2 regulation generates endocrine resistance in breast cancer. Proceedings of the National Academy of Sciences, 116(25), 12452–12461.

Mundlos, S., Otto, F., Mundlos, C., Mulliken, J. B., Aylsworth, A. S., Albright, S., … Olsen, B.R. (1997). Mutations involving the transcription factor CBFA1 cause cleidocranial dysplasia. Cell, 89(5), 773–779. doi:10.1016/s0092-8674(00)80260-3

Nagano, T., Nakashima, A., Onishi, K., Kawai, K., Awai, Y., Kinugasa, M., … Kamada, S. (2017). Proline dehydrogenase promotes senescence through the generation of reactive oxygen species. J Cell Sci, 130(8), 1413–1420. doi:10.1242/jcs.196469

Nakashima, K., Zhou, X., Kunkel, G., Zhang, Z., Deng, J. M., Behringer, R. R., & de Crombrugghe, B. (2002). The novel zinc finger-containing transcription factor osterix is required for osteoblast differentiation and bone formation. Cell, 108(1), 17–29. doi:10.1016/s0092-8674(01)00622-5

Otto, F., Thornell, A. P., Crompton, T., Denzel, A., Gilmour, K. C., Rosewell, I. R., … Owen, M.J. (1997). Cbfa1, a candidate gene for cleidocranial dysplasia syndrome, is essential for osteoblast differentiation and bone development. Cell, 89(5), 765–771. doi:10.1016/s0092-8674(00)80259-7

Phang, J. M. (2019). Proline Metabolism in Cell Regulation and Cancer Biology: Recent Advances and Hypotheses. Antioxid Redox Signal, 30(4), 635–649. doi:10.1089/ars.2017.7350

Phang, J. M., Liu, W., Hancock, C., & Christian, K. J. (2012). The proline regulatory axis and cancer. Front Oncol, 2, 60. doi:10.3389/fonc.2012.00060

Rached, M. T., Kode, A., Xu, L., Yoshikawa, Y., Paik, J. H., Depinho, R. A., & Kousteni, S. (2010). FoxO1 is a positive regulator of bone formation by favoring protein synthesis and resistance to oxidative stress in osteoblasts. Cell Metab, 11(2), 147–160. doi:10.1016/j.cmet.2010.01.001

Rodda, S. J., & McMahon, A. P. (2006). Distinct roles for Hedgehog and canonical Wnt signaling in specification, differentiation and maintenance of osteoblast progenitors. Development, 133(16), 3231–3244. doi:10.1242/dev.02480

Saikia, M., Wang, X., Mao, Y., Wan, J., Pan, T., & Qian, S. B. (2016). Codon optimality controls differential mRNA translation during amino acid starvation. Rna, 22(11), 1719–1727. doi:10.1261/rna.058180.116

Salhotra, A., Shah, H. N., Levi, B., & Longaker, M. T. (2020). Mechanisms of bone development and repair. Nature Reviews Molecular Cell Biology, 21(11), 696–711. doi:10.1038/s41580-020-00279-w

Sanjana, N. E., Shalem, O., & Zhang, F. (2014). Improved vectors and genome-wide libraries for CRISPR screening. Nat Methods, 11(8), 783–784. doi:10.1038/nmeth.3047

Sharma, D., Yu, Y., Shen, L., Zhang, G.-F., & Karner, C. M. (2021). SLC1A5 provides glutamine and asparagine necessary for bone development in mice. eLife, 10, e71595. doi:10.7554/eLife.71595

Shen, L., & Karner, C. M. (2021). Radiolabeled Amino Acid Uptake Assays in Primary Bone Cells and Bone Explants. In M. J. Hilton (Ed.), Skeletal Development and Repair: Methods and Protocols (pp. 449–456). New York, NY: Springer US.

Shen, L., Sharma, D., Yu, Y., Long, F., & Karner, C. M. (2021). Biphasic regulation of glutamine consumption by WNT during osteoblast differentiation. Journal of Cell Science, 134(1). doi:10.1242/jcs.251645

Stegen, S., Devignes, C. S., Torrekens, S., Van Looveren, R., Carmeliet, P., & Carmeliet, G. (2020). Glutamine Metabolism in Osteoprogenitors Is Required for Bone Mass Accrual and PTH-Induced Bone Anabolism in Male Mice. J Bone Miner Res. doi:10.1002/jbmr.4219

Takarada, T., Nakazato, R., Tsuchikane, A., Fujikawa, K., Iezaki, T., Yoneda, Y., & Hinoi, E. (2016). Genetic analysis of Runx2 function during intramembranous ossification. Development, 143(2), 211–218. doi:10.1242/dev.128793

Teichmann, L., Chen, C., Hoffmann, T., Smits, S. H. J., Schmitt, L., & Bremer, E. (2017). From substrate specificity to promiscuity: hybrid ABC transporters for osmoprotectants. Mol Microbiol, 104(5), 761–780. doi:10.1111/mmi.13660

Wei, J., Shimazu, J., Makinistoglu, M. P., Maurizi, A., Kajimura, D., Zong, H., … Karsenty, G. (2015). Glucose Uptake and Runx2 Synergize to Orchestrate Osteoblast Differentiation and Bone Formation. Cell, 161(7), 1576–1591. doi:10.1016/j.cell.2015.05.029

Wu, H., Whitfield, T. W., Gordon, J. A. R., Dobson, J. R., Tai, P. W. L., van Wijnen, A. J., … Lian, J.B. (2014). Genomic occupancy of Runx2 with global expression profiling identifies a novel dimension to control of osteoblastogenesis. Genome Biology, 15(3), R52. doi:10.1186/gb-2014-15-3-r52

Yang, X., Matsuda, K., Bialek, P., Jacquot, S., Masuoka, H. C., Schinke, T., … Karsenty, G. (2004). ATF4 is a substrate of RSK2 and an essential regulator of osteoblast biology; implication for Coffin-Lowry Syndrome. Cell, 117(3), 387–398. doi:10.1016/s0092-8674(04)00344-7

Yee, J. A. (1988). Effect of parathyroid hormone on amino acid transport by cultured neonatal mouse calvarial bone cells. J Bone Miner Res, 3(2), 211–218. doi:10.1002/jbmr.5650030214

Yu, Y., Newman, H., Shen, L., Sharma, D., Hu, G., Mirando, A. J., … Karner, C.M. (2019). Glutamine Metabolism Regulates Proliferation and Lineage Allocation in Skeletal Stem Cells. Cell Metab, 29(4), 966–978 e964. doi:10.1016/j.cmet.2019.01.016

Zhang, A. X., Yu, W. H., Ma, B. F., Yu, X. B., Mao, F. F., Liu, W., … Xiang, A.P. (2007). Proteomic identification of differently expressed proteins responsible for osteoblast differentiation from human mesenchymal stem cells. Mol Cell Biochem, 304(1-2), 167–179. doi:10.1007/s11010-007-9497-3

Zhang, S., Xiao, Z., Luo, J., He, N., Mahlios, J., & Quarles, L. D. (2009). Dose-Dependent Effects of Runx2 on Bone Development. Journal of Bone and Mineral Research, 24(11), 1889–1904. doi:https://doi.org/10.1359/jbmr.090502

